# Inferring the genetic basis of sleep states in *Drosophila melanogaster* using hidden Markov models

**DOI:** 10.64898/2026.01.14.699526

**Authors:** Arijit Ghosh, Susan T. Harbison

## Abstract

Multiple lines of evidence suggest that sleep in flies is not a unitary state but has lighter and deeper stages similar to mammals. A hidden Markov model applied to activity count data revealed lighter and deeper sleep states in a wild-derived population of flies. Fitting the data to a range of possible sleep states enabled us to determine the optimal number of sleep states for each fly. Four sleep and waking states were optimal across the population. We verified physiological differences among states experimentally using an arousal threshold paradigm. We then calculated the time spent in each sleep state for each fly. The time spent in each sleep state was heritable and therefore mappable to the genome. We mapped these parameters to the genome, identifying state-specific genes. Additionally, we provide software, *FlyDreamR*, that users can apply to activity count data.

## Introduction

Distinct physiological stages are a feature of sleep in mammals and humans. Polysomnography metrics enable the division of sleep into rapid-eye movement (REM) and non-rapid-eye-movement (NREM) sleep (Carskadon, 2025). Depending on how it is defined, NREM can be further divided into three to four substages of sleep (Carskadon, 2025). In general, animals and humans spend more time in NREM than in REM, with the general pattern of sleep occurring as NREM, then REM, then wake (Carskadon, 2025). Notably, the duration of each distinct sleep stage is heritable in humans. Using polysomnography in twins, evidence for a genetic component to the time spent in sleep stage has been found for all NREM stages and for REM (Ambrosius et al., 2008; Hori, 1986; Kuna et al., 2012; Linkowski et al., 1991; Linkowski et al., 1989; Webb & Campbell, 1983). Furthermore, power spectra in the 8-16 Hz range of NREM were found to have a 95% heritability (De Gennaro et al., 2008). Like humans, the time spent in paradoxical (or REM-like) sleep in mice has been shown to have a heritable component, as has the time spent in slow-wave sleep (Franken et al., 2001; Friedmann, 1974; Valatx et al., 1972). These observations suggest that multiple stages are a common feature of sleep; that they have heritable parameters that can be mapped to the genome; and that they are conserved across mammals.

Findings using disparate methodologies have converged on the idea that more than one sleep state also exists in flies. Early studies used mechanical stimuli to probe arousal threshold under the assumption that sleeping flies would be less responsive to the stimuli than waking flies (Hendricks et al., 2000a; Huber et al., 2004; Shaw et al., 2000a). These experiments established that immobile flies had an increased arousal threshold, a critical characteristic of sleep (Campbell & Tobler, 1984). Stimulating immobile flies periodically across the day revealed striking patterns: the number of flies responding to the stimulus was dependent upon the amount of time that they had been immobile and whether the stimulus was applied during the day or night (Faville et al., 2015b; van Alphen et al., 2013). Flies were less responsive after 5-10 minutes of immobility, and they were less responsive at night, suggesting that periods of immobility could be divided into different physiological states. Similar observations were made using light, air puffs, and infrared heat to stimulate the flies (Joyce et al., 2024; Keles et al., 2025; Nitz et al., 2002; Wiggin et al., 2020). Thus, quiescent flies had variable levels of arousal when immobile.

Concurrent with studies of arousal threshold were measures of physiology and electrophysiology in waking and sleeping flies. Like the arousal threshold measures, researchers found that local field potential (LFP) measures of neural activity also varied with the time spent immobile and phase (van Alphen et al., 2013; Yap et al., 2017). Specifically, LFP activity decreased in sleeping flies. Further, a genetically-encoded voltage indicator (ArcLight) and GCaMP6f calcium imaging suggested the presence of slow waves and a wake-like quiescent state, respectively, in flies (Raccuglia et al., 2019; Tainton-Heap et al., 2021). In addition, metabolic rate also decreased with time spent immobile, reaching a nadir after 35 minutes of immobility (Stahl et al., 2017). These changes in physiological measures in quiescent flies support the idea that multiple sleep states are present.

Improved videography enabled the detection of microbehaviors in flies during bouts of quiescence (Jagannathan et al., 2024; Keles et al., 2025; van Alphen et al., 2021; van Alphen et al., 2013). These microbehaviors included proboscis extension, antenna movement, and haltere switching (Jagannathan et al., 2024; Keles et al., 2025; van Alphen et al., 2021; van Alphen et al., 2013). Like the arousal threshold and neurophysiological measures, these microbehaviors varied depending on the time spent immobile and whether the measure was made during the day or night. Increased proboscis extension, periodic antennal movements, and haltere drooping mapped to periods of reduced arousal, suggesting that they are markers of deeper sleep in flies (Jagannathan et al., 2024; Keles et al., 2025; van Alphen et al., 2021; van Alphen et al., 2013). These behavioral, neuronal, and physiological measures all point to the existence of multiple sleep states in flies.

The presence of multiple sleep states is at odds with sleep derived from activity data, which is the customary method of measuring sleep in flies. Using activity data, researchers historically defined sleep as any period 5 minutes or longer without an activity count (Faville et al., 2015b; Huber et al., 2004; Nitz et al., 2002; Shaw et al., 2000a; Yap et al., 2017). Under this definition, sleep is interpreted as a unitary state, and the question arose as to whether additional information could be mined from the activity count data. Wiggin *et al*. found that it was possible to identify additional sleep and waking states within activity count data with the application of a Hidden Markov Model (HMM) (Wiggin et al., 2020). They also verified the computationally-derived sleep and wake states with arousal threshold measures using light taps. We wondered whether we could refine, optimize and apply HMMs to a large source of activity count data from a wild-derived population and derive sleep state parameters that were heritable. Doing so would enable us to derive gene networks affecting each sleep state by mapping them to the genome.

Here we applied the HMM to previously acquired sleep and activity data from 10,130 flies of the *Drosophila* Genetic Reference Panel (DGRP). Combining the HMM with this large data set enabled us to survey from 2-10 possible sleep/wake states and identify an optimal number of putative sleep states in flies. We also address the inherent uncertainties in the HMM model and provide guidelines for the optimal number of iterations needed to reach a robust readout of the state sequence for each fly. Arousal threshold measures in 28 randomly-selected genotypes demonstrate that the states predicted by the HMM correspond to physiologically lighter and deeper sleep states. We then calculate the amount of time spent in each sleep state and its heritability and use the available genomic data from the DGRP to identify candidate genes specific to each state. Last, we provide an open-source software, *FlyDreamR*, which enables the user to estimate sequence state from activity count data, to calculate the time spent in each state, and to plot the results.

## Results

### A Hidden Markov model of fly sleep

Hidden Markov models (HMMs) are a class of statistical models that represent a system as a sequence of hidden (unobserved) states linked by transition probabilities. Each hidden state stochastically generates an observable output according to a probability distribution. By assuming only the current state affects the next one (the Markov property), HMMs provide a tractable means to compute the likelihood of an observed data sequence and uncover the most probable hidden state sequence responsible for producing it. In this experiment, activity counts per minute was the observed data sequence. What we wanted to determine from this observed data was the sequence and duration of putative sleep states. *Fig. 1A* shows the key steps in the modeling process.

**Figure 1.**
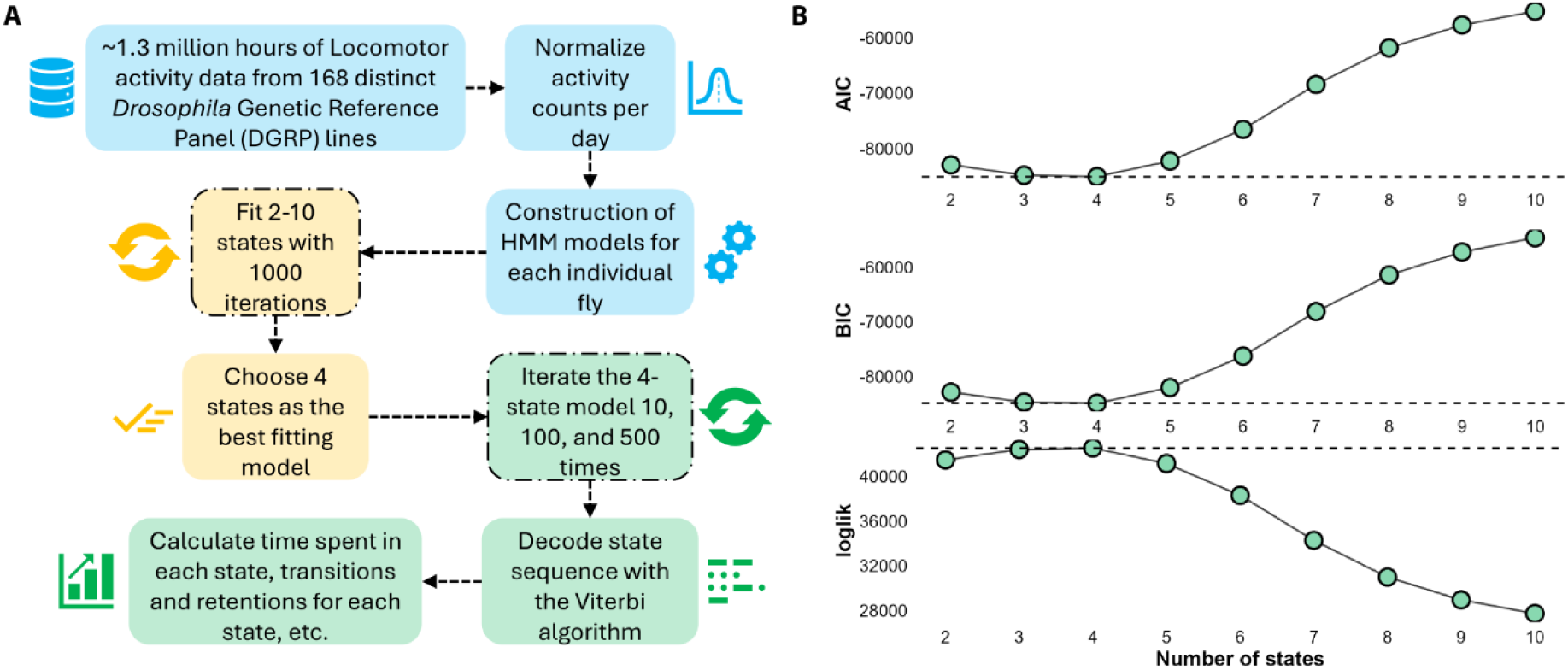
**Flowchart of analyses and rationale for the use of the 4-state model as the optimum number of sleep/wake states**. **(A)** A flowchart depicting the logic of initial analyses. **(B)** Mean Akaike and Bayesian Information Criteria (AIC and BIC), and log likelihood (loglik) values (± SE) for 2 to 10 sleep/wake states. The dashed line in each panel shows the contrast between the 4-state model and the other models. The 4-state model had the best fit to the DGRP data. The error bars are ±SE, but not visible due to their extremely small values.

#### Finding the number of putative sleep states with the best fit

A key input parameter of Hidden Markov Models (HMMs) is the number of putative “hidden” states that are assumed. We expected that the number of hidden states assumed would correspond to physiologically relevant states (Rabiner, 1989). However, evidence from previous work suggested that as few as two and as many as five wake/sleep states are detectable in flies (Jagannathan et al., 2024; van Alphen et al., 2013; Wiggin et al., 2020). Thus, the choice of the appropriate number of hidden states is not straightforward. Rather than making an assumption about the number of states that would apply to our data, we used an unbiased approach to select the optimum number of putative hidden states. We systematically examined the fit of 2 to 10 hidden states to each fly, and calculated the Akaike Information Criterion (AIC), Bayesian Information Criterion (BIC), and log likelihood (loglik) to evaluate each fit. The 4-state model had the lowest overall average AIC and BIC values (-85,053.02 and -84,931.75, respectively) and the highest overall average loglik value (42,549.51), indicating the best fit to the data (*Fig. 1B*). The 3-state model fit nearly as well, with AIC and BIC values of -84,821.27 and -84,747.45, respectively, and a loglik of 42,424.63. (*Fig. 1B*). Models with 2 or 5 states were also plausible, though they did not fit the data as well as the 4-state and 3-state models.

To further explore the appropriate number of sleep states for the HMM model, we used the BIC values for each fly separately and plotted the number of states that gave the best fit to each fly for each genotype. Depending on the fly, the lowest BIC values corresponded to 2, 3, 4, or 5 states. The 4-state model had the best fit to the majority of the flies, 39.8% overall (*Supplementary Fig. 1*). The 3-state model also fit the flies well; 30.5% flies had this model as the best fit. There were smaller numbers of flies that had the best fit to the 2-state or the 5-state model (4.4% and 25.3%, respectively). The results show that some ambiguity exists in the total number of states that are observable within a genotype.

However, our purpose was to conduct a GWAS on the time spent in each putative sleep state. In order to accomplish this, the data needed to be directly comparable from fly to fly. Given the overall best fit and plurality of flies mapping to the 4-state model, we assumed four hidden states for all subsequent calculations, therefore.

We define the four putative states as State0, State1, State2, and State3. State0 is active waking, while State3 represents the deepest sleep state. It is not entirely clear how to define States 1 and 2, and the number and defining characteristics of each sleep state is currently an unresolved question among *Drosophila* researchers. Here we will follow the convention proposed in Wiggin *et al*., that State1 represents a quiescent wake state and State2 represents light sleep, though we note that other interpretations such as a pre-sleep or active sleep state would be equally applicable (Jagannathan et al., 2024; Tainton-Heap et al., 2021; Wiggin et al., 2020).

#### Optimum number of iterations for HMM fitting and resource allocation

It is possible for the HMM to fit local rather than global maxima (C. B. Do & S. Batzoglou, 2008). This means that the sequence of putative sleep states for a given fly can vary each time the model is applied. We therefore took several steps to increase the confidence in our results. First, using the 4-state model, we applied the HMM 1000 times to the data for each fly to generate a consensus sequence. Second, we varied the initial state probabilities at each iteration by randomly drawing them from a Dirichlet distribution (Visser & Speekenbrink, 2010), a strategy that reduces any potential influence of the initial probabilities on the end state sequence. Third, we formulated an ambiguity score to quantify the degree of uncertainty in the results (see *Methods*). Using 1000 iterations per fly, we observed an ambiguity of only 0.13% across the DGRP, which suggests that the consensus state sequences we derived were robust.

However, there is a trade-off between the amount of time it takes to iterate the model repeatedly and the ambiguity of the results. Running 1000 iterations of the model on a standard desktop computer takes 3,060 ± 705 seconds per 32 flies on average (i.e., one DAM2 monitor with four days of activity and sleep counts). Thus, it may not be practical to perform many iterations on a large data set without high performance computing support. We wished to determine whether there was a minimal number of iterations that would also correspond to a minimal level of ambiguity. We ran the HMM using 10, 100, and 500 iterations and compared the results to those obtained with 1000 iterations (*Fig. 2* and *Supplementary Mov. 1*). We found that 100 iterations had low ambiguity, 0.53%, and the data were strongly correlated with 1000 iterations (*r* = ∼1, for all four states) (*Fig. 2*). On average, 100 iterations take only 7.32% as much time as 1000 iterations, 224 ± 65 seconds for 32 flies. In contrast, with 10 iterations, the correlations drop precipitously compared to 1000 iterations (*r* = 0.49 for State2 and *r* = 0.88 for State3). Thus, a minimum of 100 iterations results in unambiguous sleep state sequences while simultaneously keeping the run time relatively short.

**Figure 2.**
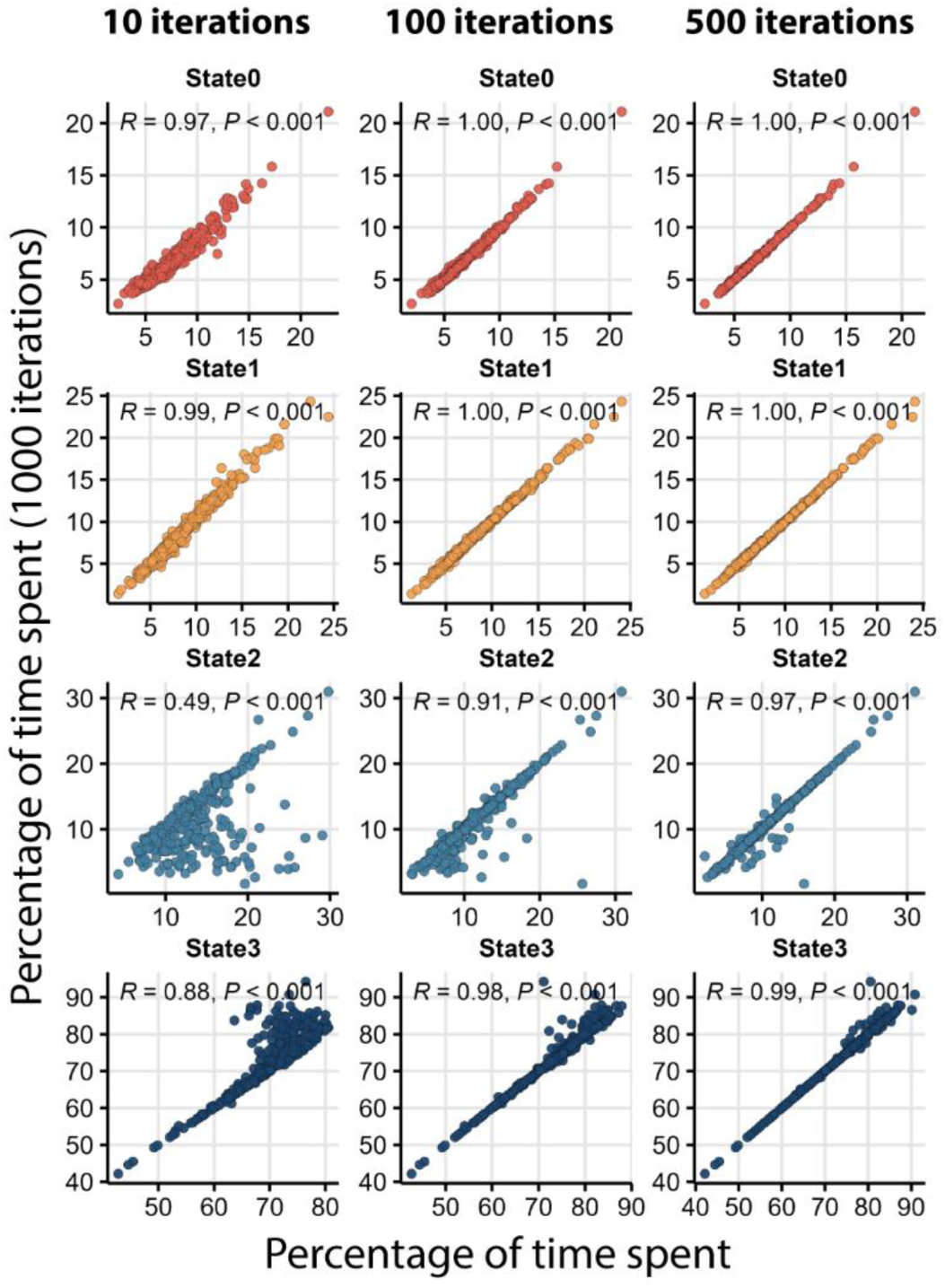
100 iterations of the HMM produce results comparable to 1000 iterations. The *y*-axis of all panels in the figure are the percentage of time spent in each state estimated using 1000 iterations. The *x*-axis represents the percentage of time spent in each state estimated using 10, 100, or 500 iterations as marked. Values used for calculating the correlations are averaged over all individuals for each genotype and sex. Pearson correlation coefficients (*R*) and significance of the correlations (*P*) are presented in each panel.

#### Experimental validation of HMM-inferred sleep states

The HMM model provided us with an output sequence of sleep states and multiple iterations enabled us to generate a consensus sequence. However, the relationship between the calculated sleep state and the actual state of the flies remained unknown. To determine whether the calculated sleep states correspond with the actual status of the animals, we probed flies from 28 randomly selected DGRP genotypes for their arousal threshold responses at different time points. We used the DART system (Faville et al., 2015a) to apply a vibration stimulus at gradually increasing levels of intensity once per hour for two continuous days (*Fig. 3A and 3B*). The DART system records fly movement using videography.

**Figure 3.**
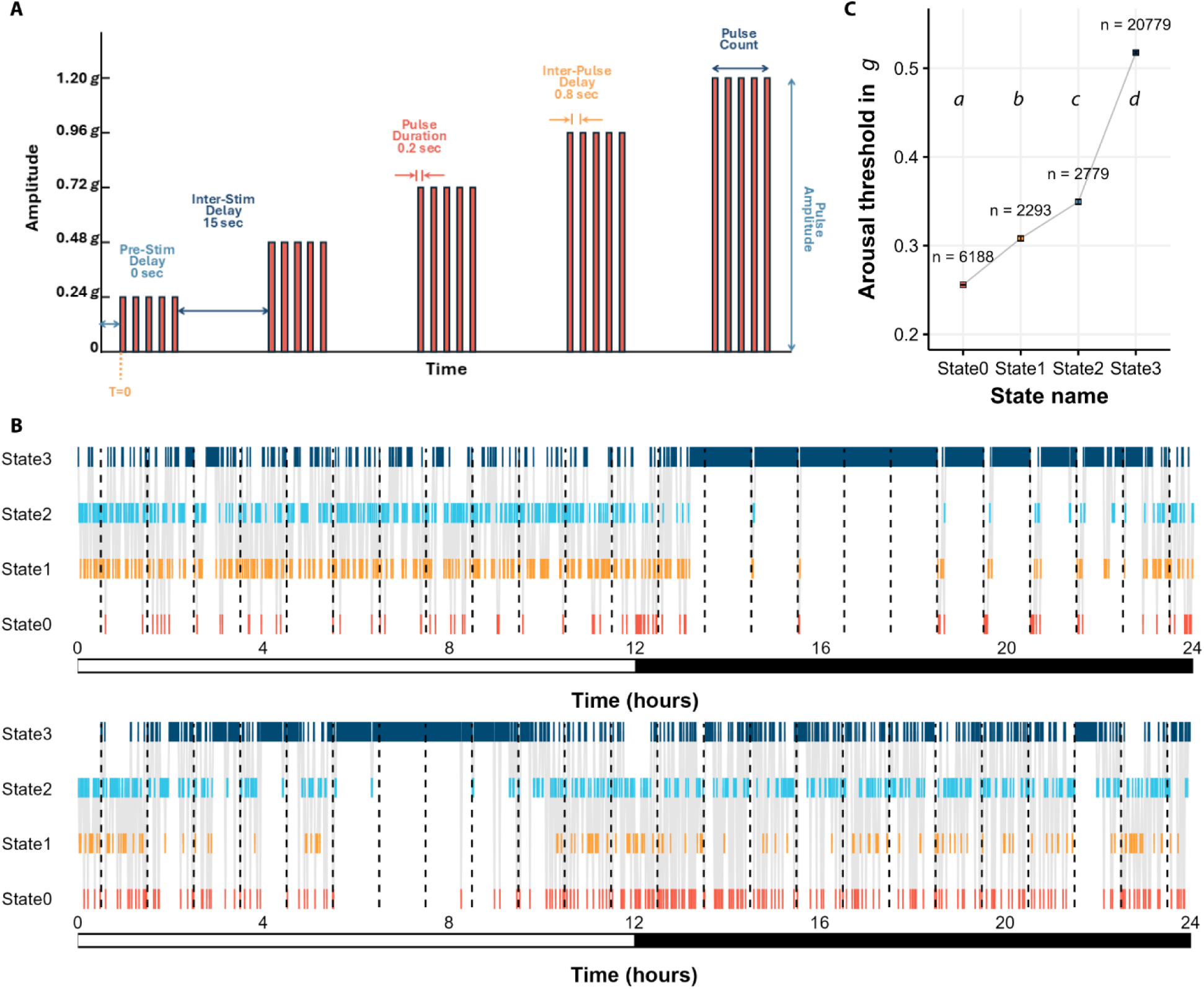
Arousal threshold probing of 28 DGRP lines detects four distinct sleep and waking states. **(A)** Schematic of 5 incremental vibration stimuli, ranging from 0.24 *g* to 1.20 *g*. The experimental parameters and order of the stimuli are shown. **(B)** Representative hypnograms of one individual fly during 2 days of arousal threshold probing. The dark dashed lines indicate times when the first stimulus in the train was applied. **(C)** Mean arousal threshold values for flies predicted to be in each sleep/wake state using the HMM. The error bars are ± SE. Mean values were significantly different from one another as indicated by letters (*P* < 0.05 and post-hoc Dunn’s analysis).

We analyzed the videos recorded in DART in two different ways: 1) absolute location tracking; and 2) tracking of midline crossing. We used the absolute location tracking data to calculate arousal threshold as was done previously (Faville et al., 2015a). Briefly, for a fly to be included in arousal threshold analysis, the fly had to be immobile before the first stimulus was applied. To determine arousal thresholds, we measured the minimum stimulus intensity that prompted resting animals to begin moving, defined as covering at least one typical body length (3mm) along the tube. We used the stimulus intensity as a proxy for the arousal threshold (van Alphen et al., 2013). If a fly did not move even after the strongest stimulus, it was marked non-respondent and was assigned a value of the highest stimulus.

The midline crossing data assessed the number of times the fly crossed a virtual midline of the tube in one-minute bins. This data is analogous to the data recorded with the DAMs. We ran the HMM model on the midline crossing data to determine which of the four states the fly was in when the vibration stimulus was applied. We then computed the average arousal threshold for each assigned sleep state.

We observed significant differences in arousal threshold among HMM-derived sleep states (*P* < 0.05). State3, the deepest sleep state, had the highest average arousal threshold (0.52 ± 0.1 *g*). The second highest arousal threshold was State2 (0.35 ± 0.06 *g*), followed by State1 (0.31 ± 0.05 *g*). State0 (0.26 ± 0.02 *g*), which is the full awake state, had the lowest arousal threshold (*Fig. 3C*). These experiments suggest that the sleep/wake states we identified from the application of HMMs on locomotor activity data correspond to physiologically different levels of arousal in the fly. We also found state-specific differences in arousal threshold in females, males, daytime, and nighttime, separately (*Supplementary Fig. 2*), demonstrating that different physiological states occur for both sexes and during any phase of the day.

### Quantitative genetic analyses of HMM-derived sleep/wake state parameters

#### Time spent in different sleep/wake states

We used the Viterbi-decoded sequences of sleep/wake states to calculate how much time the flies spent in each of the four states. We calculated the time spent in each state for day and night separately, and over 24 hours. At night, flies spent an average of 49.07 ± 0.79 minutes in State0, 36.86 ± 0.75 minutes in State1, 36.87 ± 0.85 minutes in State2, and 598.2 ± 2.06 minutes in State3. During the day, the flies averaged 51.18 ± 0.55 minutes in State0, 89.5 ± 1.13 minutes in State1, 110.92 ± 1.44 minutes in State2, and 467.65 ± 2.55 minutes in State3. Across 24 hours, flies spent an average of 100.34 ± 0.89 minutes in State0, 126.49 ± 1.5 minutes in State1, 147.96 ± 2.02 minutes in State2, and 1065.45 ± 3.65 minutes in State3 (*Fig. 4*). Thus, flies spent most of their time in the deepest putative sleep state. Flies spent more time in the lighter sleep states during the day, and more time in the deepest state at night (All *P_Phase_* <0.0001).

**Figure 4.**
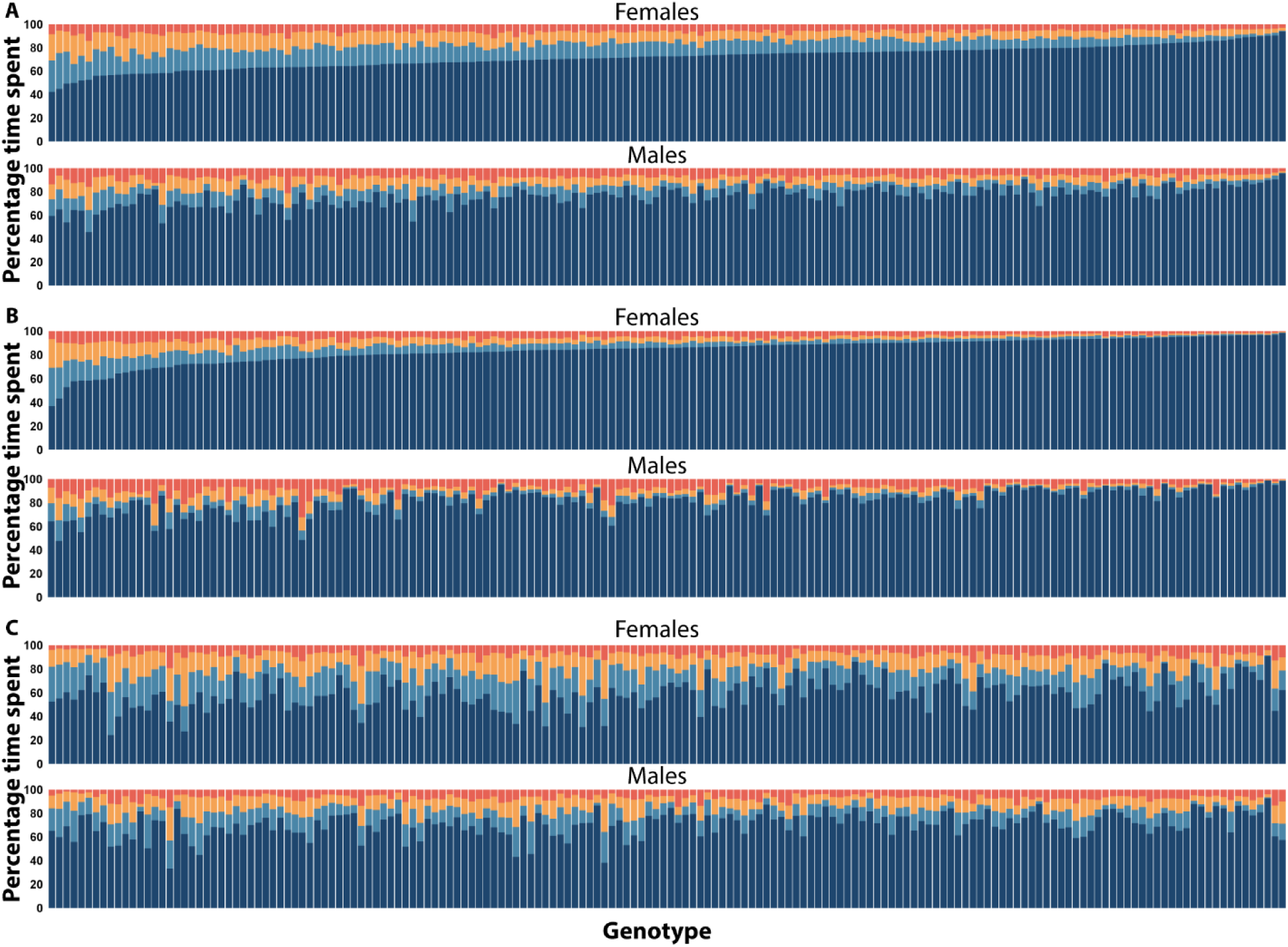
Time spent in each sleep state is genetically variable and sexually dimorphic. Mean percentage of time spent in each state plotted for females (top) and males (bottom) in **(A)** 24 hours, **(B)** the night, and **(C)** the day. All 168 genotypes are on the *x*-axis. *X*-axes are shared between **B** and **C** (i.e., each bar represents the same genotype).

The time spent in each sleep state exhibited significant genetic variation in the DGRP, for both sexes combined and for each sex separately (All *P_Line(Block)_* <0.0001). As an example, representative hypnograms for flies with long and short State3 duration are plotted in *Fig. 5*. Additionally, we observed a significant *Line×Sex* effect for all traits (All *P*_Line×Sex(Block)_ <0.0001). Broad-sense heritabilities (*H*^2^) were moderate, ranging from 0.27 – 0.53 for both sexes combined, indicating that these traits can be mapped to the genome. Sexes-separate heritability values were comparable to the sexes combined analysis. Cross-sex genetic correlations (*r*_MF_) ranged from 0.30 – 0.38; based on these relatively low values we predict that the mapping of the time spent in a given state will include sex-specific genomic variants.

**Figure 5.**
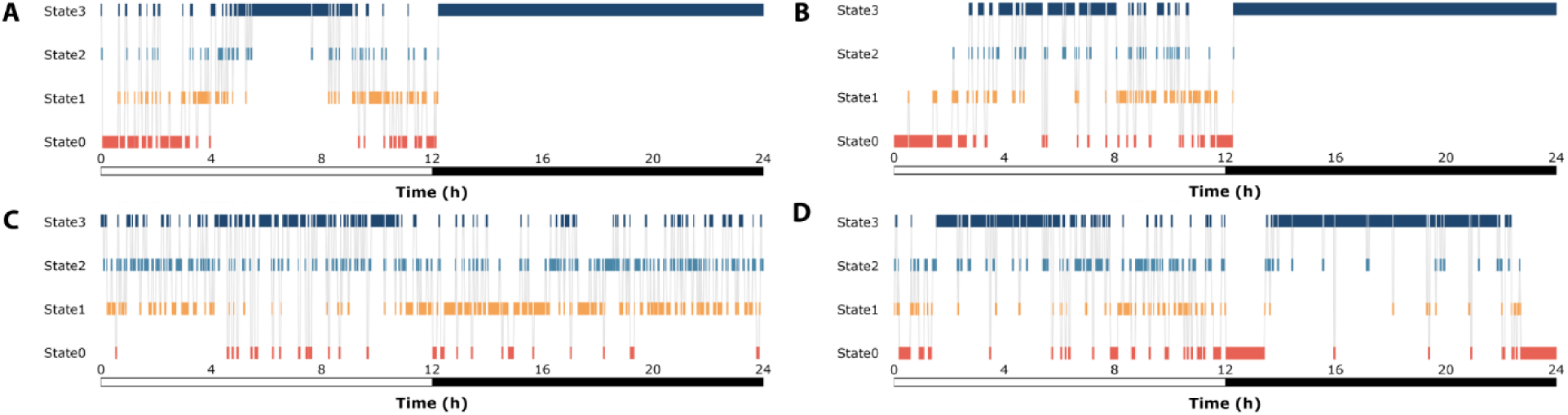
Representative hypnograms show the variety of sleep patterns in individual flies. Hypnograms show the time spent in each of the 4 sleep/wake states for each minute of the day. Time spent in each individual state is shown by vertical bars in state-specific colors. The grey lines depict transitions from one state to another. The *y*-axis depicts Zeitgeber time, and the bar below the *x*-axis depicts the light:dark cycle, where white is day, and black is night. **(A)** A representative hypnogram of a female from DGRP-41, the genotype with longest time spent in State3 at night. **(B)** A representative hypnogram of a male from DGRP-41. **(C)** A representative hypnogram of a female from DGRP-832, the genotype with the shortest time spent in State3 at night. **(D)** A representative hypnogram of a male from DGRP-832.

#### Correlations among time spent in each sleep state

We calculated phenotypic and genetic correlations (*Supplementary Fig. 3*) for time spent in the four sleep states. For nighttime sleep, correlations were statistically significant and very high. Notably, there was a strong negative correlation between the time spent in State3 and any other state. The remaining correlations were positive. Phenotypic and genetic correlations among time spent in sleep state during the day were similar to nighttime values, except for correlations between time spent in State0 and State2, which were not significant. These patterns suggest that there will be genes unique to each state, as well as genes that overlap two or more states.

In general, for a given state, the correlations between the day and night values, while statistically significant in most cases, were lower. The time spent in State0 during the day had the lowest phenotypic and genetic correlation with the time spent in State0 during the night, -0.18 and -0.11, respectively. From these values we surmised that the genetic architecture between the time spent in night sleep states would be different from the time spent in day sleep states. This result prompted us to perform the genome-wide association analyses separately for time spent in day and night in addition to the 24-hour values, for all states.

### Associations of genotype and phenotype

#### Genome-Wide Association of polymorphisms to time spent in sleep/wake states

We associated the time in each putative sleep state to 2,189,692 polymorphisms having a 5% minor allele frequency or greater in the DGRP (Huang et al., 2014). We used a genome-wide significance threshold of 1×10^-5^ (*see Methods*) (Huang et al., 2014), and identified 1,295 unique polymorphisms overall (*Fig. 6A* and *B*). Close inspection of the Q-Q plots of *P* values from the GWASs (*Supplementary Fig. 4*) also reinforces the 1×10^-5^ threshold, as do other DGRP studies (Arya et al., 2015; Dembeck, Boroczky, et al., 2015; Dembeck, Huang, et al., 2015; Eiman et al., 2024; Gardeux et al., 2023; Garlapow et al., 2015; Guzman et al., 2021; Huang et al., 2014; Hunter et al., 2016; Lobell et al., 2017; Shorter et al., 2015; Vonesch et al., 2016; Zwarts et al., 2015). Some effect sizes were very large, accounting for up to 97 minutes difference between lines having the major versus minor allele. As anticipated from studies of other complex traits using the DGRP, the distribution of effects were exponential: significant polymorphisms with higher effect sizes had lower minor allele frequencies (MAFs), while for polymorphisms with lower effect sizes, the opposite tended to be true (*Supplementary Fig. 5*) (Mackay & Huang, 2018). This pattern held whether the analysis was conducted for both sexes combined or separately; whether the polymorphism was within or outside the coding region of a gene; and whether the polymorphism was predicted to have low or moderate effects on regulation of the genes. The plot of significant polymorphisms across the genome in *Fig. 6A and B* suggested some enrichment of sleep state SNPs in particular genomic regions. Using 1Mb windows, we identified 60 genomic regions significantly enriched for sleep-state duration SNPs (*Supplementary Fig. 6*). Polymorphisms associated with time spent in 24 hours in the wake-related states (State0 and State1) showed more significantly enriched regions, particularly at the telomere end of chromosome *2L*, while polymorphisms associated with time spent in the night in the sleep-related states (State2 and State3) appeared to have more significant enriched clusters across all the major chromosomal arms, often overlapping with each other, and other states (*Supplementary Fig. 6*). Despite overlapping regions, sleep-state specific SNPs were observed.

**Figure 6.**
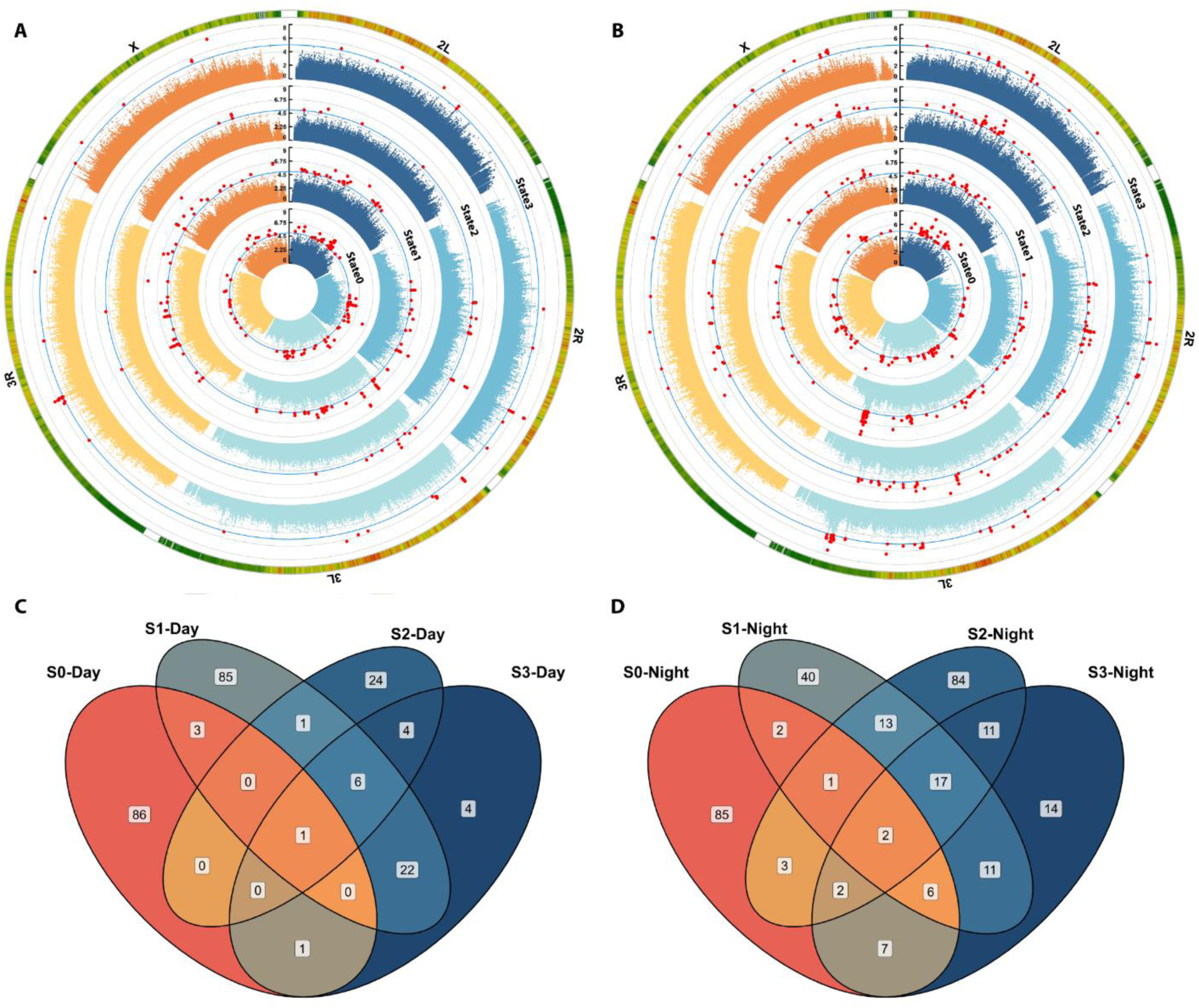
Genome-Wide Association identifies unique and common genes for time spent in sleep/wake states. **A** and **B** are circular Manhattan plots depicting significantly associated polymorphisms to time spent in 4 sleep/wake states during the **(A)** day and **(B)** night. The 5 major chromosomal arms are shown in different colors. The blue line indicates the significance threshold (1×10^-5^) of the GWAS, and the red points are the significant polymorphisms. The outermost circle is a polymorphism density heatmap, binned in 1×10^6^ base pairs. The phenotypes shown from inner to outer most circles are State0, State1, State2, and State3. **C** and **D** are Venn diagrams showing the overlap of candidate genes identified for time spent in the 4 sleep/wake states during the **(C)** day and **(D)** night.

We also examined whether presence or absence of *Wolbachia* infection and chromosomal inversions associated with sleep states, as local LD increases in regions of inversions. We found that time spent in State0 in 24 hours and daytime is associated with *In_3R_Mo* and *In_2L_t*, respectively; time spent in State1 in 24 hour and daytime is associated with *In_2R_NS* and *In_2L_t*, respectively, and time spent in State3 in the daytime is associated with *In_2L_t*. Some polymorphisms associated with time spent in distinct sleep/wake states fall within these inversions (see below) and are therefore confounded with them.

#### Identification of significantly associated polymorphisms and genes

We identified 505 unique genes within 1kb of 989 polymorphisms. Genes were often unique to a given sleep state (*Fig. 6C and D*). The vast majority of the polymorphisms identified were SNPs (79.7%), but there were some deletions (11.1%), insertions (9.07%), and mixed polymorphisms (0.06%). Most of the polymorphisms were within introns (67.22%); followed by variants falling inside transcripts (10.5%); and downstream, intergenic, and upstream variants (8.76%, 7.12%, and 6.38%, respectively). 73 variants were predicted to have a moderate effect on protein structure and thus function, including missense variants, conservative and disruptive in-frame insertions, and deletions. Of the 505 genes identified, 376 had human orthologs (Hu et al., 2011).

#### Types of genes influencing the time spent in sleep state

Certain themes emerged through our study of these genes. Many genes had roles in neurotransmission; known or predicted enzymatic activity; roles in well-known signaling pathways; functions in transcription or were transcription factors; or were involved in the structural components of the cell, such as the extracellular matrix, extracellular region, or cytoskeleton.

### Neurotransmission

We saw several associations with genes having roles in neurotransmission. Notably, the *5-HT2B* serotonin receptor was associated with the time spent in State3 during the day, a gene known to affect sleep time and rebound after sleep deprivation (Qian et al., 2017). Also, *sNPF-R* was associated with the time spent in State0 at night and was previously implicated in sleep (Chen et al., 2013). Additionally, *Proc-R,* a gene implicated in sleep latency, was associated with the time spent in State1 during the day (Eiman et al., 2024). This theme was reinforced by genes modulating neurotransmission across multiple states. Examples include the nicotinic acetylcholine receptor subunit *nAChRalpha3* and *qvr*, the latter regulating cholinergic transmission and associated with nearly all nighttime sleep states (State 1/2/3-Night) (Koh et al., 2008; Pimentel et al., 2016; Wu et al., 2010). The vesicular monoamine transporter, *Vmat*, was linked to all four states, highlighting the global role of neurotransmitters (e.g., dopamine, serotonin) in modulating sleep (Harbison et al., 2013; Nall & Sehgal, 2013). Thus, neurotransmitters have roles unique to the time spent in a particular sleep state as well as functioning across sleep states.

### Enzymatic activity

We noted that roughly 20% of the genes we identified had either known or predicted enzymatic activity (103 genes out of 505). Oxidoreductases comprise 14% of the enzymes in the fly genome; transferases, 33%; hydrolases, 37%; lyases, 3%; isomerases, 7%; ligases, 3%; and translocases, 3%. Within our list, however, hydrolases were over-represented (46%) and transferases were under-represented (24%) (Both *P* <0.001, hypergeometric test). Hydrolases are enzymes that use a water molecule to break a chemical bond, suggesting that time spent in sleep state may have a role in the breakdown of complex molecules. Hydrolases were associated with the time spent in all sleep states during the day and night but were a particularly prominent feature of the time spent in State2 at night (15 out of 23 enzymes). These State2 hydrolases have known or predicted roles in the breakdown of ATP, triacylglycerols, and chitin; in the DNA damage response; and in proteolysis, among other functions (Ozturk-Colak et al., 2024). As mentioned earlier, there were fewer transferases than would be expected. Notably, three of the four enzymes mapping to the time spent in State2 during the day were kinases. The presence of many varied enzymes on our gene lists suggests that their activity is an important feature underlying the time spent in each sleep state.

### Pathways

Flybase tracks genes that affect 17 signaling pathways, including pathways previously implicated in fly sleep such as EGFR, Insulin-like receptor, Notch, and Wnt signaling pathways (Foltenyi et al., 2007; Harbison et al., 2013; Harbison et al., 2017; Metaxakis et al., 2014; Seugnet et al., 2011). Core genes functioning in each pathway as well as genes known to be positive or negative regulators of the pathway are tracked. Genes for night sleep states map to 14 of these pathways, but the number of genes is not enriched for any specific pathway or sleep state. Many of the genes we mapped are known to be positive or negative regulators of these pathways; however, there were some genes that were core pathway members. Core genes for the BMP, EGFR, and Wnt pathways mapped to the time spent in State2 during the night and day . Core genes for the Toll receptor pathway mapped to the time spent in State0 during the night and day, respectively.

### Transcription factors

We found that 9.7% of the genes associated with time spent in sleep state were transcription factors, cofactors, or had roles in mRNA splicing. These included *foxo*, a transcription factor that impacts sleep during the day in flies with reduced insulin signaling (Metaxakis et al., 2014). Nineteen genes were identified in a GWAS of sleep summary phenotypes previously (Eiman et al., 2024; Harbison et al., 2019; Harbison et al., 2013; Wu et al., 2018). The presence of transcription factors could indicate a signal cascade relevant to sleep initiation or maintenance, a hypothesis we test below.

### Human homologs

Additionally, some identified genes have human homologs known to affect various traits, e.g. cocaine addiction, circadian entrainment, Alzheimer’s disease, neurodegenerative diseases, highlighting the evolutionary conservation of sleep and related mechanisms and potential relevance of discoveries made if flies for understanding mammalian or human sleep regulation (Kanehisa et al., 2016).

#### Gene ontology and network analysis

A gene ontology analysis revealed that time spent in the deepest sleep state in day and night are enriched in genes associated with learning, memory, axon and neuronal development, and immune responses and motor neuron axonal guidance, respectively; while the waking states in day and night are enriched in genes associated with detection and sensory perception of temperature stimulus, and negative regulation of axonogenesis and response to light stimulus, respectively. In total, we found 1,130 GO terms to be significantly enriched (FDR ≤ 0.05).

Genetic and protein interaction network analysis showed a significant enrichment in protein-protein interactions for State0 (night and 24-hour), State1 (day and night), State2 (night), and State3 (night) genes (*Fig. 7*). These networks enabled us to identify certain “hub” genes which are highly connected with other genes in the network. Importantly, we identified hub genes which are common across day and night traits, and may control time spent in different sleep/wake states throughout the day and night, albeit by different mechanisms. We found that 17 of these genes overlapped with 254 unique genes known or predicted to affect different aspects of sleep and circadian rhythm (Singh et al., 2024). Prior GWAS data on sleep shows an overlap of 245 genes out of 2,168 previously observed (Eiman et al., 2024; Harbison et al., 2019; Harbison et al., 2013; Wu et al., 2018).

**Figure 7.**
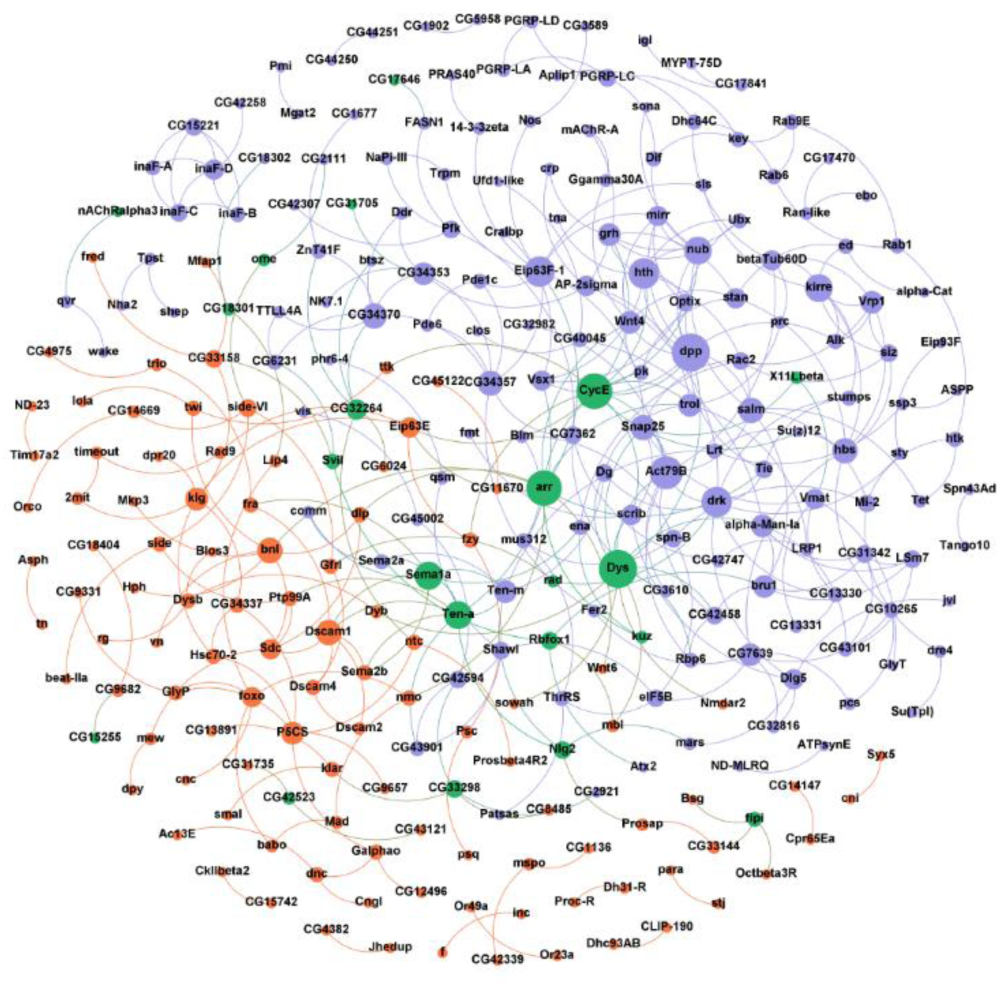
Interaction network of genes reveals ‘hub’ genes. The figure shows a STRING interaction network of the genes/proteins associated with time spent in 4 sleep/wake states in day (orange nodes), night (purple nodes), and common among both day and night (green nodes). The node sizes depict the degree of the node. Unconnected nodes were removed from the final network.

## Discussion

Here we find evidence for multiple sleep states using a computationally-derived model to interpret activity data from 10,130 flies. Application of the HMM is feasible on a standard desktop computer with only 100 iterations.

### More than one sleep state exists in natural populations of Drosophila

As we noted in the Introduction, orthogonal measures of arousal threshold, microbehavioral movements, physiology, and neuronal electrical activity suggest that more than one sleep state is present in flies (Faville et al., 2015b; Jagannathan et al., 2024; Joyce et al., 2024; Keles et al., 2025; Nitz et al., 2002; Raccuglia et al., 2019; Stahl et al., 2017; Tainton-Heap et al., 2021; van Alphen et al., 2021; van Alphen et al., 2013; Wiggin et al., 2020; Yap et al., 2017). Difficulty in translating these findings across studies emerges as most measures of sleep made in flies are based on activity count data, such as readings made using the Trikinetics *Drosophila* Activity Monitor (DAM). Our application of the HMM to a large dataset of activity count data from the DGRP found statistical support for two to five sleep/wake states in individual flies. We observed some ambiguity in the optimal number of states within a genotype, as well as variation in the number of states among genotypes. This could be a *bona-fide* biological phenomenon, or it could be a limitation of the HMM’s ability to distinguish among states. Historically, the description of sleep in humans faced similar challenges. An early account of sleep and waking in humans described five levels of sleep based on EEG measures (Loomis et al., 1936). This was further refined with the distinction between REM and NREM observed by Dement and Kleitman in 1957 (Dement & Kleitman, 1957). Later work proposed that NREM be divided into four stages (N1, N2, N3, and N4) (Rechtschaffen, 1968), until the modern definition of sleep by the American Academy of Sleep Medicine proposed combining the deepest two stages, N3 and N4, into one (Iber, 2007). Differences in the amount of time assigned to each sleep stage in a particular individual occur depending on which standard is applied (Moser et al., 2009). Thus, ambiguity in determining the appropriate number of sleep states is not unique to flies.

Four sleep/wake states were the most prevalent in our data, followed by three states. Using the HMM, Wiggin *et al*. also uncovered evidence for four sleep/wake states in several genotypes, which they termed full wake, early wake, light sleep, and deep sleep (Wiggin et al., 2020). In contrast, detection of sleep states using other methods found support for three and five sleep/wake states. Sleep deprivation employing a yoked control only showed effective perturbation of the sleep homeostat when sleep bouts 25 minutes or longer were disrupted, implying three overall sleep/wake states: a lighter sleep or quiescent waking state and a deeper sleep state along with an active waking state (Chowdhury et al., 2023). Three states are also supported by a two-process model analysis showing that long sleep bouts (30 minutes or greater) are the most important contributors to the relationship between circadian period and sleep amount (Abhilash & Shafer, 2024). Another study combining electrophysiology with microbehavioral measures found that five overall sleep/wake states could be defined (Jagannathan et al., 2024). Interestingly, two states, which correspond to one mode of waking and one mode of sleep, were the least prevalent in our data: it was the best fit model for only 4.4% of the flies. This suggests that consideration of these additional states is required to fully understand sleep and its genetic underpinnings in flies.

#### Robust classifications of sleep state emerge from activity count data with relatively few iterations

Computational methods such as the HMM rely on iterations that converge to a solution. It is possible for individual runs of the model to produce different state sequences as the model can converge on local rather than global maxima (Chuong B. Do & Serafim Batzoglou, 2008). Thus, the HMM should be run multiple times on the data for each fly to account for these potential differences. However, it is not clear how many runs are needed to produce reliable data. We explored the robustness of state classification by varying the number of iterations of the HMM fitting process. Notably, we found that 100 iterations were sufficient to produce state computation time with minimal loss of accuracy. We note that the model did not converge for a small subset of fly activity data (1.1%). Why the model did not converge in these cases is not clear as we did not detect any trend among these flies, though we anecdotally observed longer run times with long-sleeping flies. One possibility is that very long sleep bouts, which correspond to strings of zeros in the activity count data, provide less information for the HMM to interpret. Nevertheless, our analysis demonstrates that a moderate number of iterations can adequately infer sleep/wake states, which is beneficial for scaling up to large datasets. Additionally, a standard laptop/desktop is sufficient to run the HMM model for small datasets.

#### HMM-inferred sleep/wake states correspond to different physiological states

Although repeated iterations of the HMM increased our confidence in the model’s predictions, we lack a “ground truth” dataset to compare our sleep/wake sequences to—and this is the case for measuring sleep in any organism. However, sleep-like states have been associated with elevated arousal threshold in response to subtle mechanical stimuli, compared to awake flies. Responsiveness to stimuli decreases from lighter to deeper states of sleep (Faville et al., 2015a; Hendricks et al., 2000b; Jagannathan et al., 2024; Joyce et al., 2024; Keles et al., 2025; Nitz et al., 2002; Shaw et al., 2000b; Tainton-Heap et al., 2021; van Alphen et al., 2013; Wiggin et al., 2020; Yap et al., 2017). To validate our HMM results, we used a video monitoring system to track both activity and arousal responses to a gradually increasing mechanical stimulus simultaneously. From this data we generated activity counts analogous to the data produced by the Trikinetics DAM system by simulating an infrared beam across the fly activity monitor tubes and analyzing the data using the HMM. We found that a distinct arousal threshold maps to each sleep and wake state predicted by the HMM. Flies in the putative deep sleep state (State3) were least likely to respond even with the highest intensity stimulus. Those in the light sleep state (State2) had a lower arousal threshold, followed by the quiescent wake (State1) state. The fully awake state (State0) had the lowest arousal threshold. Our data reinforce earlier studies using tapping (Wiggin et al., 2020), or frequent air puffs (Joyce et al., 2024) to explore the predictions of the HMM. Our validation experiment suggests that the sleep/wake states inferred by the HMM are indeed physiologically distinct and not just arbitrary mathematical constructs. Importantly, the model applies to the highly variable sleep seen among wildtype flies of the DGRP (Harbison et al., 2013). The HMM applied to activity monitoring can therefore serve as a proxy to effectively assess sleep depth in lieu of electrophysiology data, enabling the continued use of activity monitoring for large-scale genetic screens and genome mapping. A future goal is to determine whether candidate genes identified using activity count methods contribute to changes in sleep state observed using other methods such as electrophysiology or video analysis of microbehaviors.

#### Genetic associations of time spent in sleep/wake states reveal state-specific genes

Here we show that quantification of different characteristics of sleep states, such as the time spent in each sleep state, is also feasible with our refinement of the HMM. We demonstrated that the time spent in each state exhibited moderate broad-sense heritability, suggesting that these parameters could be mapped to the genome. Genome-wide association identified 1,295 unique polymorphisms associated with time spent in specific states across 24 hours, day, and night. Mapping significant variants to the genome (±1 kb) identified 505 genes. Most variants were intronic (∼67%), with 73 variants predicted to have moderate coding defects. These results extend the prior GWAS on total sleep amount (Harbison et al., 2013) by demonstrating that sleep micro-architecture, the allocation of time among sleep/wake states, is itself a genetically regulated complex trait. Our findings indicate that sleep intensity (depth) and sleep quantity are at least partially separable at the genetic level. Some loci specifically bias time in deep sleep, light sleep, or quiet wake, while others act across states, cohering with the genetic correlation structure.

#### Biological and conceptual implications

First, demonstrating multi-state sleep from locomotor records (validated physiologically via arousal measurement) supports evolutionary conservation of sleep architecture. Second, enrichment of sensory-detection pathways among wake-state genes and neuronal development/learning among deep-sleep genes suggests distinct molecular levers for sensory gating vs restorative depth, consistent with the notion that animals tune sleep intensity to environmental safety and context. Third, circadian modulation of state allocation and state-specific associations indicates that the clock not only times sleep but helps set its quality (depth distribution) across the day.

Additionally, our approach could be adapted to other behaviors and species. The HMM’s success in parsing sleep indicates potential for classifying sub-states of other complex behaviors like feeding (e.g. distinguishing sipping vs. proboscis grooming vs. long feeding bouts) or gait patterns in locomotion (McClintock et al., 2020). Applying this framework to different *Drosophila* species and across taxa one can compare sleep evolution (Joyce et al., 2024). Studying candidate hub genes through network-level perturbation could reveal how single disruptions affect the sleep-regulatory network. Finally, integrating these GWAS hits with time-of-day transcriptomics/proteomics could illuminate how circadian outputs gate entry into, and retention of, deep vs light sleep.

## Conclusion

We have developed an HMM-based approach to analyze *Drosophila* sleep/wake behavior, identifying multiple reproducible states akin to mammalian sleep stages. This approach allowed detailed sleep quantification and uncovered robust correlations with both physiology (arousal threshold) and genotype. Using the DGRP, we mapped the genetic architecture influencing sleep states, identifying genes and networks related to sleep depth. Combining this approach with functional genomics and neurobiology will help link genes and neuronal populations to specific sleep states. By mapping sleep states, we hope to explore why animals, from flies to humans, need periods of quiescence and how evolution balances wakefulness and rest.

## Materials and Methods Description of the data used

We utilized a previously collected dataset from our lab (Harbison et al., 2013). 168 *Drosophila* Genetic Reference Panel (DGRP) lines were assayed for 7 days under a 12-hour light-dark cycle (LD12:12) at 25°C with standard corn meal food. The lines were randomly divided into four blocks, with each block assayed in four replicates, resulting in measurements from 32 virgin flies per line per sex, totaling 10,239 flies.

We analyzed four days of data for each individual fly, excluding the first day of recording. The total data analyzed comprised ∼1.3 million hours of locomotor activity recorded using the *Drosophila* Activity Monitoring (DAM) system (Trikinetics, Waltham, MA), encompassing 10,130 individual flies from 168 unique genotypes. Data from 109 flies were not used due to reasons explained in the *Fitting the HMMs* section.

## HMM construction and fitting

One of the key initial decisions is how many “hidden” states are to be fitted. Rather than assume an arbitrary number of hidden states, we used an unbiased approach. We fitted HMMs with 2 to 10 putative sleep/wake states to the data for each individual fly to determine the optimal number of states. The best performing model was chosen based on Akaike and Bayesian Information Criteria (AIC and BIC) (Akaike, 1974; Schwarz, 1978) and log likelihood values (Costa & De Angelis, 2010; Rabiner & Juang, 1986; Rabiner, 1989). The four-state model had the best fit according to these parameters (see Results for details). We labeled each of the four states as State0, State1, State2, and State3, with State0 representing the most active state (highest activity counts) and State3 representing the deepest putative state of sleep (lowest activity counts).

The HMM models were constructed as outlined by Rabiner (Rabiner, 1989) and fitted using the depmixS4 package (Visser & Speekenbrink, 2010). This package facilitates the definition and estimation of HMMs, with parameters estimated via the expectation-maximization (EM) algorithm. The HMMs were constructed using the *depmix* function from the *depmixS4* library, specifying these transition and initial probability matrices, with the response variable being the normalized activity levels per minute for a whole day (1440 data points), and fitted with the *fit* function from the *depmixS4* library. We then used the Viterbi algorithm to decode the data, resulting in a sequence of sleep states for each fly for each day. *Fig. 1A* summarizes key steps in this process briefly.

### Data initialization and normalization

Activity counts from the DAM system fluctuate among genotypes, sexes, individuals, and days. To create a uniform standard of comparison, we divided the activity counts for each minute by the total activity for each day, then multiplied the result by 100. Activity levels thus fluctuated from 0 to 100 for each fly each day. These normalized activity levels were then used to build the HMMs.

Additionally, we defined an initial probability matrix, which captures the likelihood that the underlying Markov chain starts in a particular sleep state before any observations are made. Formally, they are a set of probabilities (usually denoted by π) that sum to 1 and describe how the process “chooses” its initial state. For the first iteration, the initial transition matrix was constructed such that transition probabilities between the two most extreme states (i.e., State0 and State3 and vice versa) were set to a small value (1×10^-05^). The remaining initial state probabilities were then set equally across all states. We then conducted as many as 999 additional iterations. For these subsequent iterations, random initial state probabilities were drawn from a Dirichlet distribution (Visser & Speekenbrink, 2010) (See *Accounting for local maxima* section for more details).

### Fitting the HMMs

Transition probabilities define how a Markov chain advances through its states at discrete time steps. They are expressed in a matrix *A*, where each entry *aᵢⱼ* specifies the probability of moving from state *i* at time *t* to state *j* at time *t+1*. The rows of *A* must each sum to *1*, denoting that the system must occupy one and only one state at each subsequent step. Observation probabilities, denoted by *B*, determine how likely it is that a particular observed activity count is generated by each state. Together with the initial probability distribution over states, these parameters (*A* and *B*) characterize the hidden Markov model’s ability to capture temporal dynamics in the data.

To estimate the hidden state probabilities, we used the Baum-Welch algorithm (Baum et al., 1970) iteratively to reach the best explanation of the observed data—in this case, the pattern of activity counts. It applies the Expectation-Maximization (EM) principle in two repeating steps. First, in the “E-step,” a forward-backward algorithm is used to estimate how likely it is for the model to be in each hidden state (and to make each transition) at every time point, under the current parameter settings. Next, in the “M-step,” the algorithm updates the model’s parameters (the initial probabilities, the probabilities of observing the given activity levels in each state, and the probabilities of switching from one state to another) such that they result in a higher log-likelihood. Subsequent cycles of these two steps push the model’s log-likelihood upward until the improvement in log-likelihood becomes very small (a relative difference of 1×10^-08^ between the latest and previous model), signaling that the model has converged. In rare instances, we noted that the models for some of the data did not converge. We removed 109 flies (1.06%) from the original dataset of 10,239 flies due to failed convergence after 100,000 trials.

### Decoding sleep state sequences

After fitting the model, state sequences were decoded using the Viterbi algorithm (Viterbi, 1967) to obtain the most likely sequence of hidden states. This algorithm determines the sequence of sleep states with the greatest likelihood of being generated by the observed activity count data. States exhibiting higher activity levels were assigned to lower-number states, while states with lower activity levels were assigned to higher-number states. For example, in the 4-state model, each state is defined as was conceptualized previously (Wiggin et al., 2020): State0 had the highest mean activity and corresponds to waking; State1 had the second highest activity and roughly corresponds to quiescent waking, State2 had the third highest activity and roughly corresponds to light sleep, and State3 had the lowest activity and corresponds to deep sleep.

#### Accounting for local maxima

The likelihood surface navigated by the EM algorithm can contain multiple local maxima. This means that each time we run the EM algorithm with different random initial values, it is possible to generate a different sequence of sleep states for the same input data. We took several steps to ensure that we obtained the optimal state sequence. To mitigate differing solutions, we conducted multiple runs of the EM algorithm and compared the resulting state sequences. We employed randomized initialization for each iteration. The EM fitting process was repeated 10, 100, 500, and 1000 times to generate many Viterbi-decoded state sequences for the same data. A consensus state sequence was then constructed by selecting the most dominant state, i.e., the one appearing most frequently at each of the 1440 time points in a day. We hypothesized that the most frequent state sequence was the best fit to the data and confirmed this later with experimental tests of arousal threshold. We defined an ambiguity score for each state sequence using the formula *(1 - (n / T)) * 100*, where *n* = number of times the most frequent state sequence appeared and *T* is the total number of iterations. Lower ambiguity scores reflect a greater confidence in the state sequences, while higher ambiguity scores would equate to lower confidence in the state sequence. We used 1000 iterations to generate all of the phenotypic data we used in the GWAS, but we also compared these results with phenotypes derived using fewer iterations (see Results for details).

### Sleep state quantification with HMMs

Following the fitting of the HMMs using the 4-state model with 1000 iterations, we quantified the time spent in each identified sleep/wake state in minutes. This analysis was conducted for the entire 24-hour period, as well as separately for day and night phases. The data were then averaged over the total number of days for each fly. These calculations and quantifications were performed in R, using custom-written codes.

### An R implementation of the HMM pipeline – the FlyDreamR package

The pipeline described above was converted into a user-friendly R package – *FlyDreamR.* The package enables users to run an HMM on sleep and activity data generated using the Trikinetics *Drosophila* Activity Monitoring (DAM) System (Trikinetics, Waltham, MA). The package is available under an MIT license. The package has three main functions – *HMMDataPrep*, *HMMbehavr* (and its parallel and much faster implementation, *HMMbehavrFast*), and *HMMplot*. The *HMMDataPrep* function prepares DAM data for use with the HMMs. HMMDataPrep binds user-supplied metadata (such as genotype, sex, block, replicate, and treatment) to DAM activity count files. From here the user can calculate sleep parameters using the traditional 5-minute definition of sleep or a user-defined definition (such as 60 minutes). The HMMbehavr function performs the primary iterative HMM fitting procedure and generates useful tables for downstream applications, including the time spent in each sleep state for day, night and 24 hours. *HMMbehavrFast* is a parallel implementation of *HMMbehavr* and can process the data more rapidly, depending on the number of processors allocated to the function. *HMMplot* enables the user to plot heatmap hypnograms for a large number of individuals over multiple days efficiently. Accessory functions such as *HMMFacetedPlot* and *HMMSinglePlot* offer visualization and options for saving large numbers of expanded hypnograms. All functions have numerous customization options and detailed help pages explaining these options and outputs. The *FlyDreamR* package is also accompanied by a GUI – the *Shiny-FlyDreamR* app, which will also make it accessible to users unfamiliar with programming to run the HMMs on their own data.

## Experimental validation of sleep states through arousal threshold

The HMM-inferred putative sleep/wake states correspond to lighter and deeper sleep states, as well as quiescent and fully awake states. To validate these inferred states, we conducted arousal threshold probing in flies. We hypothesized that flies in deeper sleep states would exhibit significantly higher arousal thresholds compared to flies in lighter sleep states or awake states. Flies were placed in glass tubes used with DAM systems and recorded via infrared videography for four days in .avi format at 5 frames per second (FPS). Arousal threshold was measured as described previously (Faville et al., 2015a) using the DART apparatus (bfklab, Hertford, UK). Briefly, shaftless vibrating motors delivered a vibrational stimulus to the flies. The vibration amplitudes are expressed by the unit *g* (gravitational force of earth at the surface; 9.8m/s^2^). The DART apparatus is controlled by a commercially available MATLAB-based software made by bfklab (bfklab, Hertford, UK). Video processing and stationary fly detection is performed with a Convoluted Neural Network (CNN) classification model, trained on videos from the same experiment, implemented with the MATLAB Deep Learning Toolbox. As a result of only recording in the infrared range, videos captured are homogenous across day and night, experiments, and not affected by changes in lighting conditions inside the incubators.

We randomly chose 28 genotypes from the 168 DGRP genotypes assayed for sleep above for the DART-based arousal threshold measurements. The lines were divided into seven blocks at random, with each block being tested in two replicates for four days under LD12:12 and 25°C. This resulted in measurements from 28 virgin flies per line per sex, totaling 1,568 flies and 150,528 hours of video recording. From the start of Day 3 of video recording, the flies were subjected to a progressively stronger vibrational stimulus train every hour, and we assessed whether they woke up immediately following each stimulus. The stimuli ranged from 0.24 *g* to 1.2 *g* in five equal steps (*Fig. 3A*). We recorded the stimulus level that elicited movement from each fly, defined as traveling 3 mm (Faville et al., 2015b). For instances where the fly did not move after receiving the stimuli, it was assigned the highest stimulus level, 1.2 *g*’s. Data from flies that were already mobile when the stimuli were applied were removed from the analysis, and data from any dead flies was removed as well. Next, we used the video recordings to calculate the number of times each fly crossed a virtual line positioned at the middle of the tube, generating data analogous to the beam crossing data produced by the DAM system. We used this simulated beam-crossing data to run HMMs and classify sleep state sequences. We sorted arousal threshold levels according to HMM-classified sleep state. We compared arousal threshold level across sleep state using a Wald test. Flies were also tracked to their absolute position and sleep measurements were calculated on both beam-crossing data and absolute location data. The HMM analyses for the beam-crossing data were performed in R. All sleep parameter calculations from the video recordings were performed in MATLAB, for both beam-crossing and absolute location tracking data. This experiment enabled us to correlate experimentally derived arousal thresholds with the corresponding computationally derived sleep state.

## Quantitative genetic analyses

We partitioned the variance in time spent in 4 sleep/wake states in 24-hour, day, and night using the general linear model *Y = μ + B + S + L(B) + S × L(B) + R(B) + S × R(B) + R × L(B) + S × R × L(B) + ε*, where *L* (Line), *R* (Replicate), and *B* (Block) are random effects, *S* (Sex) is a fixed effect, and ε is the error variance. Reduced models were used to partition variances for each sex as *Y = μ + B + L(B) + R(B) + R × L(B) + ε*. Variance components were measured using the restricted maximum likelihood (REML) method in order to compute broad-sense heritability (*H^2^*) as described previously (Harbison et al., 2013) For sexes combined, *H^2^* was calculated as *H^2^ = (σ^2^_L_ + σ^2^_SL_)/(σ^2^_L_ + σ^2^_SL_ + σ^2^_E_)*, where *σ^2^_L_* is variance component among lines, *σ^2^_SL_* is the Line × Sex variance component, and *σ^2^_E_* is all other sources of variation. For each sex separately, we used the formula *H^2^ = σ^2^_L_/(σ^2^_L_ + σ^2^_E_)* for estimating broad-sense heritability.

We calculated the genetic correlation (*r_G_*) between time spent in sleep states using *r_G_ = cov_ab_/√(σ ^2^ × σ_Lb_^2^)*, where *cov_ab_* is the covariance between metrics *a* and *b*, and *σ_La_ ^2^* and *σ_Lb_^2^* are variances among lines for metrics *a* and *b*, respectively. For each sleep state, the genetic correlation between the day and night state was statistically significant but less than 1.0 (e.g., the genetic correlation between the time spent in State0 during the day and the time spent in State0 during the night was -0.16). Thus, some differences between day sleep and night sleep exist, as has been demonstrated previously (van Alphen et al., 2013). The statistical models were conducted using SAS/STAT (15.3), and downstream analyses were performed using R.

## Associations of genotype and phenotype

We performed genome-wide association analyses for all 12 traits using the line means of each trait with 2,189,692 segregating polymorphisms in the DGRP, using the DGRP2 web-based analysis tool (https://quantgenet.msu.edu/dgrp/) (Huang et al., 2014; Mackay et al., 2012), updated according to the release 6 of *Drosophila melanogaster* genome assembly and annotation. This tool performs genotype-phenotype associations using Fast-LMM (Lippert et al., 2011). The line means are adjusted for the effects of *Wolbachia pipientis* infection status and five reported major inversions in the DGRP, *In(2L)t*, *In(2R)NS*, *In(3R)K*, *In(3R)P*, and *In(3R)Mo*, wherever present, before performing the GWAS. The webtool fits a linear mixed model of the form *y =* **X***b +* **Z***u + e*, where *y* is the line mean of a given trait, adjusted for effects of *Wolbachia* infection and inversions, **X** is the covariance relationship matrix for the fixed variant effect *b*, **Z** is the incidence matrix for the random polygenic effect *u*, and *e* is the residual effect. Effect sizes of different variants are estimated as half of the difference between line means bearing the major and minor alleles. Effect sizes were normalized by their phenotypic standard deviation. We used a genome-wide threshold *P*-value of 1×10^-5^ (Arya et al., 2015; Dembeck, Boroczky, et al., 2015; Dembeck, Huang, et al., 2015; Eiman et al., 2024; Gardeux et al., 2023; Garlapow et al., 2015; Guzman et al., 2021; Huang et al., 2014; Hunter et al., 2016; Lobell et al., 2017; Shorter et al., 2015; Vonesch et al., 2016; Zwarts et al., 2015) for selecting significant polymorphisms from the GWAS results. The webtool also provides a list of the genes located within a ±1kb distance of the significant polymorphisms. The results of the GWASs were plotted as circular Manhattan plots (*Fig. 6A & B*) with CMplot (Yin et al., 2021). The Q-Q plots of the *P*-values (*Supplementary Fig.* 4) were also plotted with CMplot (Yin et al., 2021).

## Pathway and Network analyses

All genes associated with each trait were used for gene ontology (GO) of biological processes and network analysis with ShinyGO (Ge et al., 2020) and stringApp within Cytoscape (Doncheva et al., 2023; Shannon et al., 2003; Szklarczyk et al., 2023) with default options, respectively. The networks were visualized using Gephi (Bastian et al., 2009). We also investigated if certain regions of the genome were enriched in SNPs significantly associated with each trait. To achieve this, we performed hypergeometric tests in 1Mb non-overlapping steps using the base R function *phyper*. For each window we assessed whether it contained a significantly greater number of SNPs compared to what would be by chance, given the chromosome-wide distribution of significant and non-significant SNPs. The resulting *P*-values from the hypergeometric tests were adjusted using the Benjamini-Hochberg false discovery rate (FDR) method. Over- and under-representation analysis of enzymes were performed using the base R function *phyper*, using the total number of enzymes in the *Drosophila* genome as background.

All plots were generated in R using the *ggplot2* library. All statistical analyses were performed using R, and all statistical significances are inferred at α = 0.05, unless otherwise mentioned. All variability measures in the text within parentheses are ±SD, unless otherwise mentioned.

## Supporting information

Supplementary Mov. 1

## Acknowledgements

The authors would like to thank W. Huang, R. Faville, and I. Jain for technical assistance. This work used the computational resources of the National Institutes of Health High-Performance Computing Biowulf cluster (http://hpc.nih.gov).

## Author contributions

Conceptualization: AG, STH Methodology: AG, STH Investigation: AG

Formal Analysis: AG Software: AG Resources: STH

Writing – Original Draft: AG

Writing – Review and Editing: AG, STH

## Data and materials availability

The *FlyDreamR* package and the *Shiny-FlyDreamR* app is available through GitHub (https://github.com/orijitghosh/FlyDreamR), and the published version will be archived on Zenodo (link to be provided if manuscript is published). We accessed publicly available Release 6 genome variant data for the DGRP via https://quantgenet.msu.edu/dgrp/. Additional information on the DGRP, including the SRA accession numbers for the DNA sequences of each DGRP line, can be found in Huang et al.

---

**Supplementary Figure 1.**
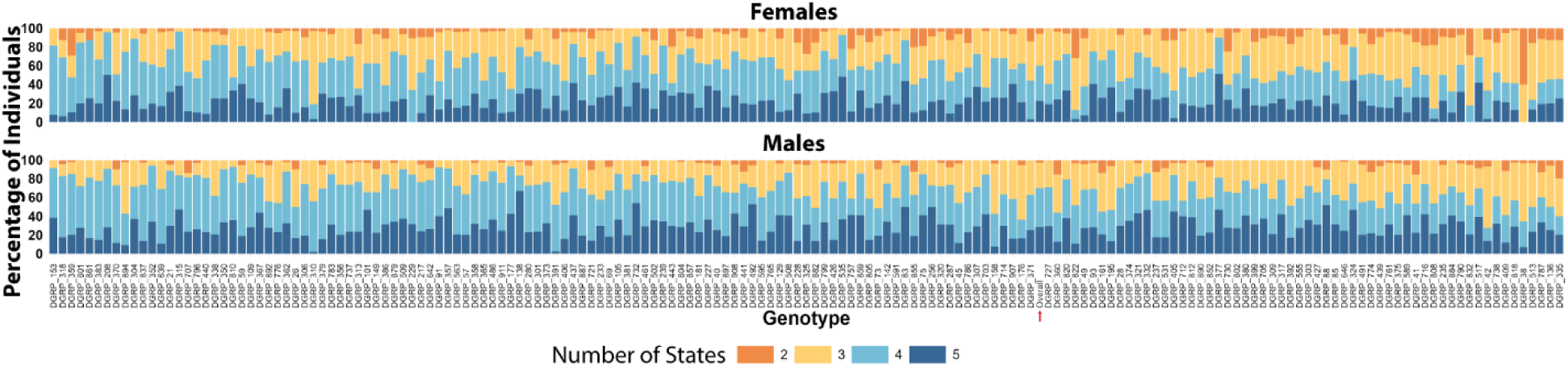
Percentage of individuals fitting 2, 3, 4, and 5 sleep/wake states across the **DGRP.** Among all 168 genotypes in the DGRP, 4.4% showed best fit to 2 states, 30.5% to 3 states, 39.8% to 4 states, and 25.3% to 5 states. Numbers of individuals indicating best fit to each number of states are represented as percentages to compare across sexes (top row females and bottom row males) and genotypes. All 168 genotypes are plotted across the *x*-axis. The “overall” on the x-axis marked with a red arrow represents the mean of all 168 genotypes.

**Supplementary Figure 2.**
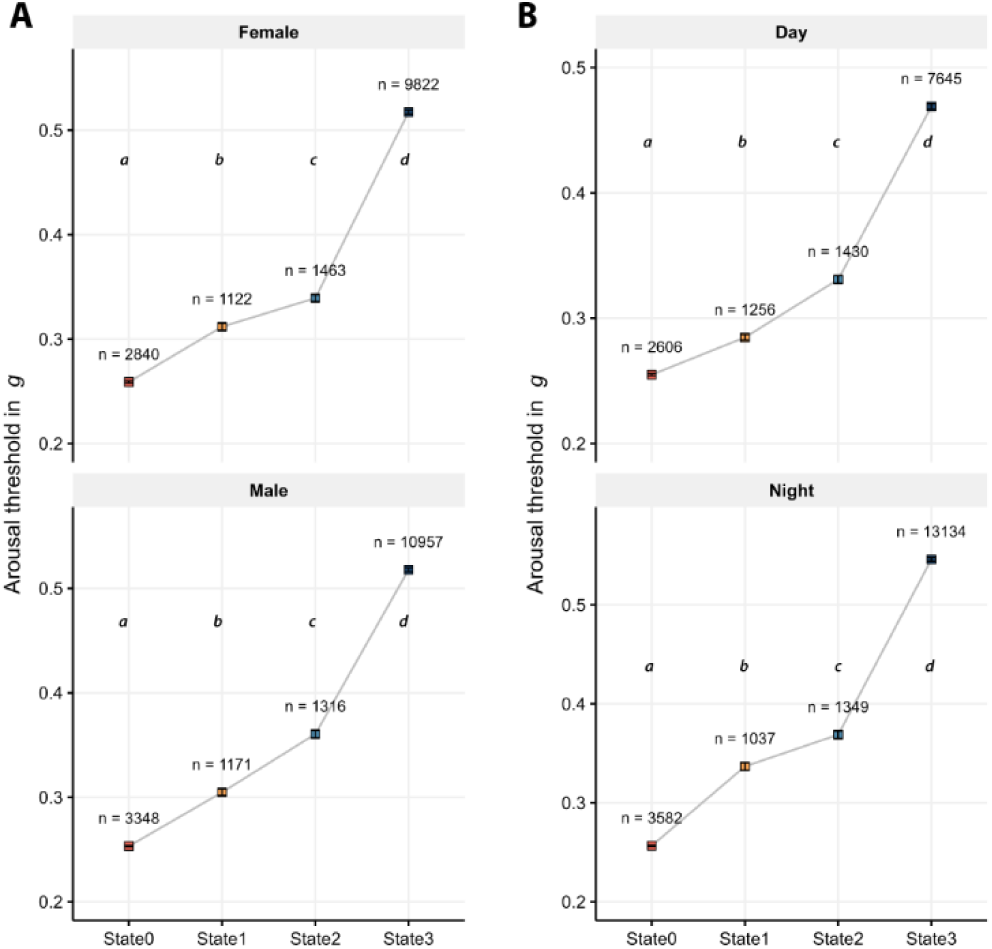
Arousal threshold probing of 28 DGRP lines detects four distinct sleep and waking states irrespective of sex and time of day. A and B are mean arousal threshold values for flies predicted to be in each sleep/wake state using the HMM in different sexes, and different times of day, respectively. The error bars are ± SE. Mean values were significantly different from one another as indicated by letters (*P* < 0.05 and post-hoc Dunn’s analysis).

**Supplementary Figure 3.**
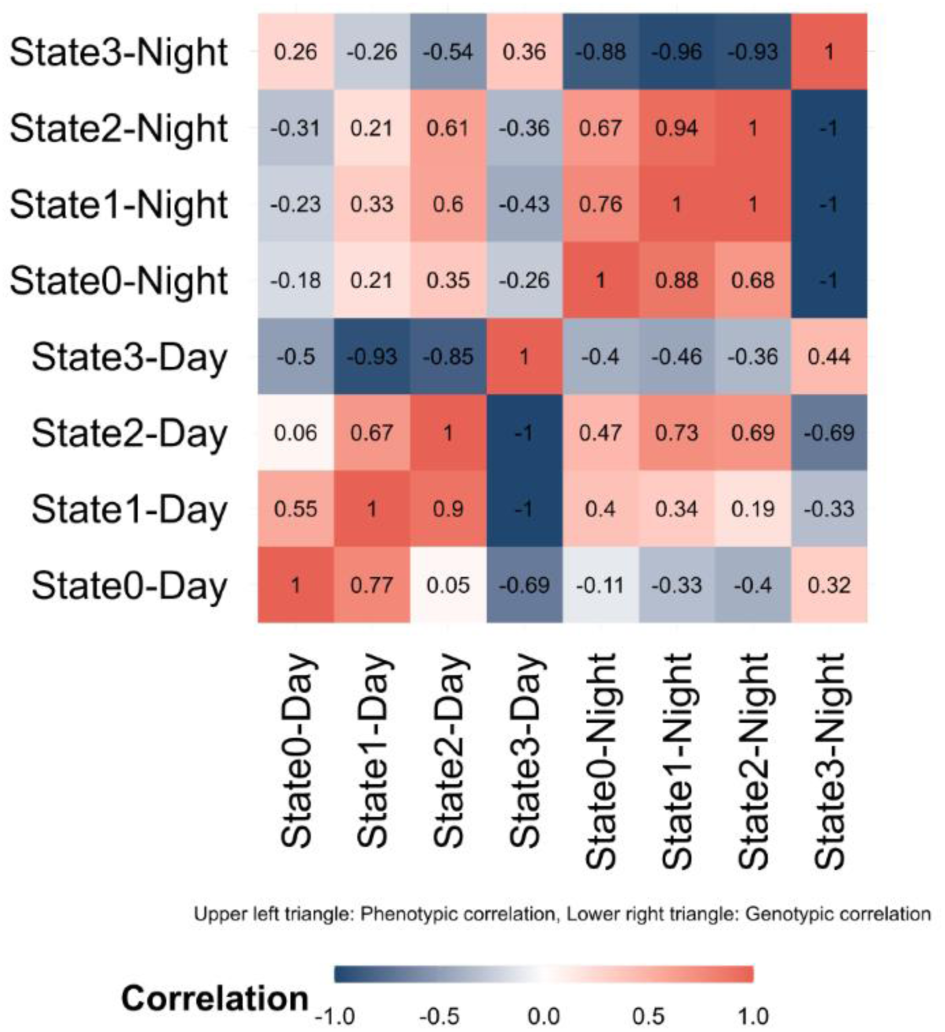
Heatmap of phenotypic and genotypic correlation coefficients among time spent in 4 sleep/wake states in day and night. The phenotypic (upper left triangle) and genotypic (lower right triangle) correlation coefficients were calculated as described in the *Materials and Methods* section. Correlation values range from +1 (red) to -1 (blue). In general, for a given state, the correlation between time spent in day and night, while statistically significant in most cases, were lower than correlations within a given phase, indicating that the genetic architecture between time spent in night states and day states is likely to be different.

**Supplementary Figure 4.**
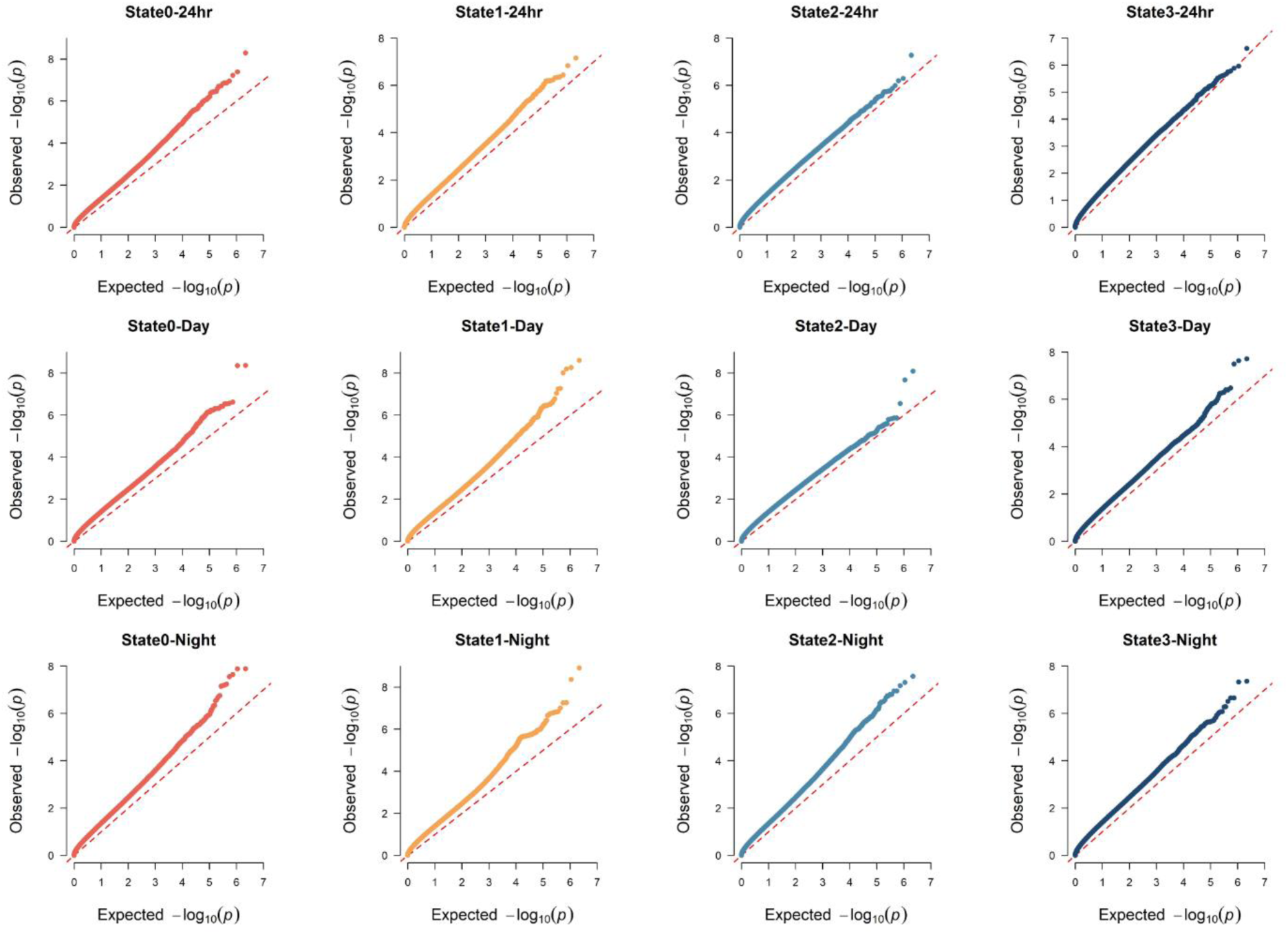
Quantile-Quantile (Q-Q) plots illustrating observed vs expected distributions of negative log-transformed *P*-values (*-log_10_(P)*) from the GWASs for each trait. The top, middle, and bottom rows show Q-Q plots for into 24-hour, daytime, and nighttime states, respectively. Each point represents a SNP tested in the analysis. The dashed red lines indicate the expected distribution under the null hypothesis, highlighting deviations indicative of significant genetic associations.

**Supplementary Figure 5.**
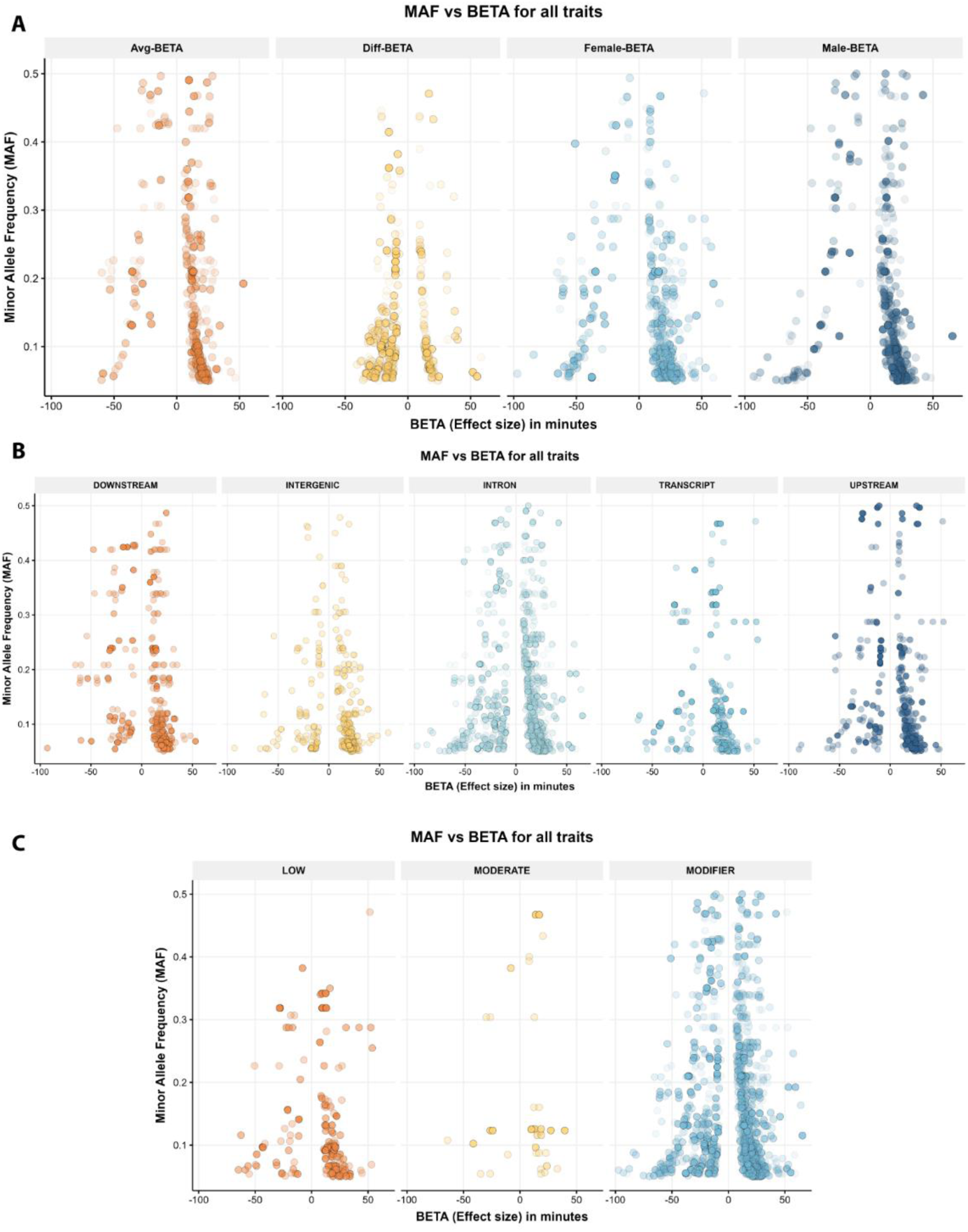
: Relationship between Minor Allele Frequency (MAF) and effect sizes (BETA, measured in minutes) for polymorphisms associated with different traits. (A) Comparison of MAF against effect sizes for average (Avg-BETA), differential (Diff-BETA), female-specific (Female-BETA), and male-specific (Male-BETA) polymorphisms. (B) Effect size distribution categorized by genomic annotations: downstream, intergenic, intron, transcript, and upstream regions. (C) Distribution of effect sizes grouped by predicted functional impact: low, moderate, and modifier categories. Each data point represents an individual polymorphism, highlighting the variation in allele frequencies and their phenotypic effects across traits and genomic contexts.

**Supplementary Figure 6:**
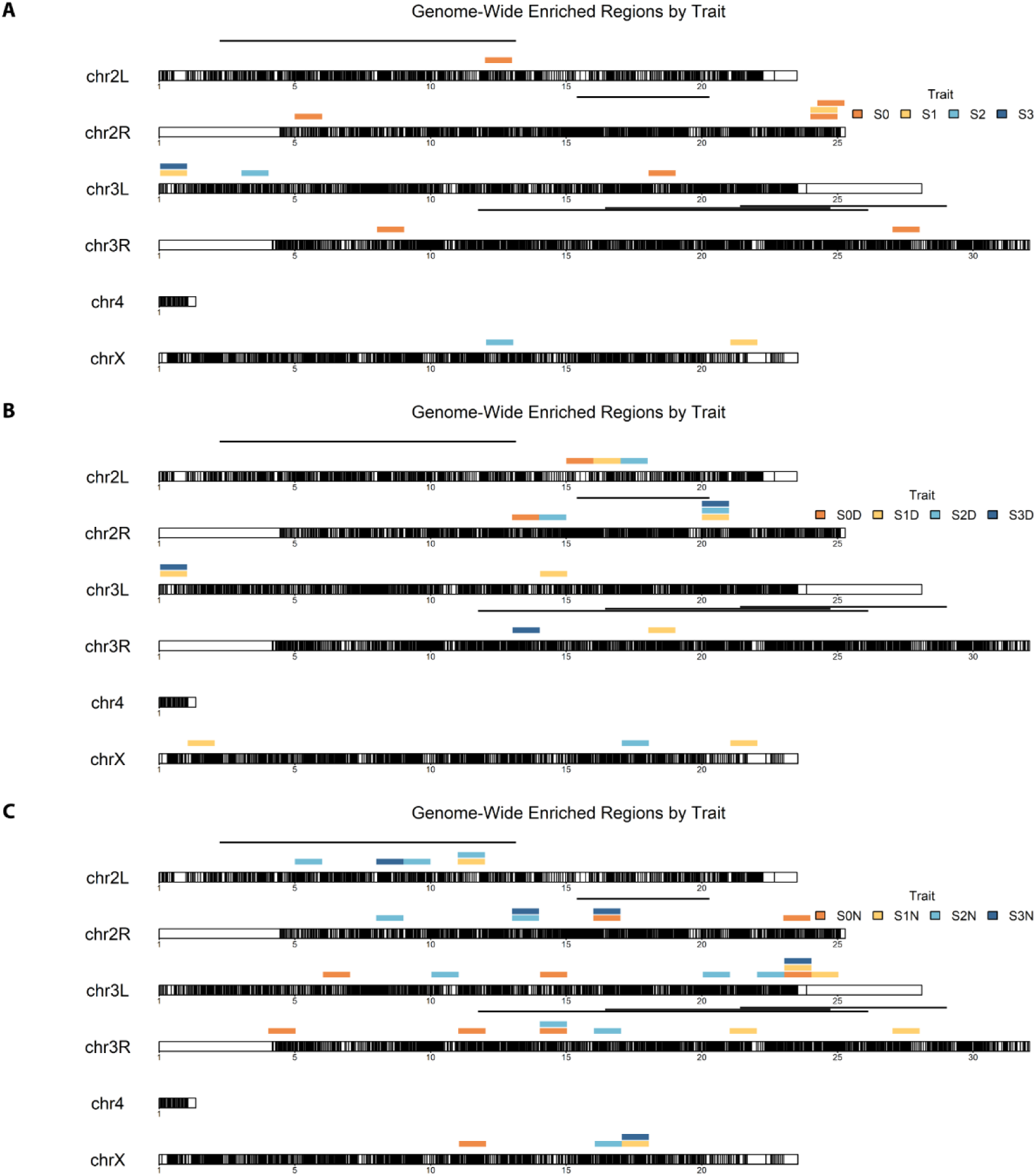
Genome-wide representation of significant SNP enriched regions across chromosomes in the DGRP. Panels depict genomic regions significantly enriched in SNPs associated with time spent in 4 sleep/wake states **(A)** in the 24-hour period, **(B)** in the day, and **(C)** in the night. Colored bars indicate a 1-Mb stretch of significant genomic regions enriched for each trait (S0: State0, S1: State1, S2: State2, and S3: State3), mapped onto chromosomal arms (*2L*, *2R*, *3L*, *3R*, *4*, and *X*). The black bars show the positions of 5 major chromosomal inversions present in the DGRP.

## References

1. Abhilash, L., & Shafer, O. T. (2024). A two-process model of Drosophila sleep reveals an inter-dependence between circadian clock speed and the rate of sleep pressure decay. Sleep, 47(2). 10.1093/sleep/zsad277

2. Akaike, H. (1974). A new look at the statistical model identification. Ieee Transactions on Automatic Control, 19(6), 716–723. 10.1109/tac.1974.1100705

3. Ambrosius, U., Lietzenmaier, S., Wehrle, R., Wichniak, A., Kalus, S., Winkelmann, J., Bettecken, T., Holsboer, F., Yassouridis, A., & Friess, E. (2008). Heritability of sleep electroencephalogram. Biol Psychiatry, 64(4), 344–348. 10.1016/j.biopsych.2008.03.002

4. Arya, G. H., Magwire, M. M., Huang, W., Serrano-Negron, Y. L., Mackay, T. F., & Anholt, R. R. (2015). The genetic basis for variation in olfactory behavior in Drosophila melanogaster. Chem Senses, 40(4), 233–243. 10.1093/chemse/bjv001

5. Ayoob, J. C., Terman, J. R., & Kolodkin, A. L. (2006). Drosophila Plexin B is a Sema-2a receptor required for axon guidance. Development, 133(11), 2125–2135. 10.1242/dev.02380

6. Baggio, F., Bozzato, A., Benna, C., Leonardi, E., Romoli, O., Cognolato, M., Tosatto, S. C., Costa, R., & Sandrelli, F. (2013). 2mit, an intronic gene of Drosophila melanogaster timeless2, is involved in behavioral plasticity. PLoS One, 8(9), e76351. 10.1371/journal.pone.0076351

7. Bastian, M., Heymann, S., & Jacomy, M. (2009). Gephi: An Open Source Software for Exploring and Manipulating Networks. Proceedings of the International AAAI Conference on Web and Social Media, 3(1), 361–362. 10.1609/icwsm.v3i1.13937

8. Baum, L. E., Petrie, T., Soules, G., & Weiss, N. (1970). A Maximization Technique Occurring in the Statistical Analysis of Probabilistic Functions of Markov Chains. The Annals of Mathematical Statistics, 41(1), 164–171, 168.

9. Campbell, S. S., & Tobler, I. (1984). Animal sleep: a review of sleep duration across phylogeny. Neurosci Biobehav Rev, 8(3), 269–300. 10.1016/0149-7634(84)90054-x

10. Carskadon, M. A. (2025). Definitions and Basic Methods for Assessing Sleep. Fundamentals of Sleep and Circadian Science, 1.

11. Chen, W., Shi, W., Li, L., Zheng, Z., Li, T., Bai, W., & Zhao, Z. (2013). Regulation of sleep by the short neuropeptide F (sNPF) in Drosophila melanogaster. Insect Biochem Mol Biol, 43(9), 809–819. 10.1016/j.ibmb.2013.06.003

12. Chowdhury, B., Abhilash, L., Ortega, A., Liu, S., & Shafer, O. (2023). Homeostatic control of deep sleep and molecular correlates of sleep pressure in Drosophila. Elife, 12. 10.7554/eLife.91355

13. Costa, M., & De Angelis, L. (2010). Model selection in hidden Markov models : a simulation study [Working Paper](2010/7). (Quaderni di Dipartimento. Serie Ricerche, Issue. https://amsacta.unibo.it/id/eprint/2909/

14. De Gennaro, L., Marzano, C., Fratello, F., Moroni, F., Pellicciari, M. C., Ferlazzo, F., Costa, S., Couyoumdjian, A., Curcio, G., Sforza, E., Malafosse, A., Finelli, L. A., Pasqualetti, P., Ferrara, M., Bertini, M., & Rossini, P. M. (2008). The electroencephalographic fingerprint of sleep is genetically determined: a twin study. Ann Neurol, 64(4), 455–460. 10.1002/ana.21434

15. Dembeck, L. M., Boroczky, K., Huang, W., Schal, C., Anholt, R. R., & Mackay, T. F. (2015). Genetic architecture of natural variation in cuticular hydrocarbon composition in Drosophila melanogaster. Elife, 4. 10.7554/eLife.09861

16. Dembeck, L. M., Huang, W., Magwire, M. M., Lawrence, F., Lyman, R. F., & Mackay, T. F. (2015). Genetic Architecture of Abdominal Pigmentation in Drosophila melanogaster. PLoS Genet, 11(5), e1005163. 10.1371/journal.pgen.1005163

17. Dement, W., & Kleitman, N. (1957). Cyclic variations in EEG during sleep and their relation to eye movements, body motility, and dreaming. Electroencephalography and Clinical Neurophysiology, 9(4), 673–690. 10.1016/0013-4694(57)90088-3

18. Dissel, S., Angadi, V., Kirszenblat, L., Suzuki, Y., Donlea, J., Klose, M., Koch, Z., English, D., Winsky-Sommerer, R., van Swinderen, B., & Shaw, P. J. (2015). Sleep restores behavioral plasticity to Drosophila mutants. Curr Biol, 25(10), 1270–1281. 10.1016/j.cub.2015.03.027

19. Do, C. B., & Batzoglou, S. (2008). What is the expectation maximization algorithm? Nature Biotechnology, 26(8), 897–899. 10.1038/nbt1406

20. Do, C. B., & Batzoglou, S. (2008). What is the expectation maximization algorithm? Nat Biotechnol, 26(8), 897–899. 10.1038/nbt1406

21. Doncheva, N. T., Morris, J. H., Holze, H., Kirsch, R., Nastou, K. C., Cuesta-Astroz, Y., Rattei, T., Szklarczyk, D., von Mering, C., & Jensen, L. J. (2023). Cytoscape stringApp 2.0: Analysis and Visualization of Heterogeneous Biological Networks. J Proteome Res, 22(2), 637–646. 10.1021/acs.jproteome.2c00651

22. Eiman, M. N., Kumar, S., Serrano Negron, Y. L., Tansey, T. R., & Harbison, S. T. (2024). Genome-wide association in Drosophila identifies a role for Piezo and Proc-R in sleep latency. Sci Rep, 14(1), 260. 10.1038/s41598-023-50552-z

23. Faville, R., Kottler, B., Goodhill, G. J., Shaw, P. J., & van Swinderen, B. (2015a). How deeply does your mutant sleep? Probing arousal to better understand sleep defects in Drosophila. Scientific Reports, 5(1). 10.1038/srep08454

24. Faville, R., Kottler, B., Goodhill, G. J., Shaw, P. J., & van Swinderen, B. (2015b). How deeply does your mutant sleep? Probing arousal to better understand sleep defects in Drosophila. Sci Rep, 5, 8454. 10.1038/srep08454

25. Foltenyi, K., Greenspan, R. J., & Newport, J. W. (2007). Activation of EGFR and ERK by rhomboid signaling regulates the consolidation and maintenance of sleep in Drosophila. Nat Neurosci, 10, 1160–1167.

26. Franken, P., Chollet, D., & Tafti, M. (2001). The homeostatic regulation of sleep need is under genetic control. J Neurosci, 21(8), 2610–2621. 10.1523/JNEUROSCI.21-08-02610.2001

27. Friedmann, J. K. (1974). A diallel analysis of the genetic underpinnings of mouse sleep. Physiol Behav, 12(2), 169–175. 10.1016/0031-9384(74)90169-3

28. Gallio, M., Ofstad, T. A., Macpherson, L. J., Wang, J. W., & Zuker, C. S. (2011). The coding of temperature in the Drosophila brain. Cell, 144(4), 614–624. 10.1016/j.cell.2011.01.028

29. Garbe, D. S., Vigderman, A. S., Moscato, E., Dove, A. E., Vecsey, C. G., Kayser, M. S., & Sehgal, A. (2016). Changes in Female Drosophila Sleep following Mating Are Mediated by SPSN-SAG Neurons. J Biol Rhythms, 31(6), 551–567. 10.1177/0748730416668048

30. Gardeux, V., Bevers, R. P. J., David, F. P. A., Rosschaert, E., Rochepeau, R., & Deplancke, B. (2023). DGRPool: A web tool leveraging harmonized Drosophila Genetic Reference Panel phenotyping data for the study of complex traits. Elife, 12. 10.7554/eLife.88981.1

31. Garlapow, M. E., Huang, W., Yarboro, M. T., Peterson, K. R., & Mackay, T. F. (2015). Quantitative Genetics of Food Intake in Drosophila melanogaster. PLoS One, 10(9), e0138129. 10.1371/journal.pone.0138129

32. Ge, S. X., Jung, D., & Yao, R. (2020). ShinyGO: a graphical gene-set enrichment tool for animals and plants. Bioinformatics, 36(8), 2628–2629. 10.1093/bioinformatics/btz931

33. Guven-Ozkan, T., Busto, G. U., Schutte, S. S., Cervantes-Sandoval, I., O’Dowd, D. K., & Davis, R. L. (2016). MiR-980 Is a Memory Suppressor MicroRNA that Regulates the Autism-Susceptibility Gene A2bp1. Cell Rep, 14(7), 1698–1709. 10.1016/j.celrep.2016.01.040

34. Guzman, R. M., Howard, Z. P., Liu, Z., Oliveira, R. D., Massa, A. T., Omsland, A., White, S. N., & Goodman, A. G. (2021). Natural genetic variation in Drosophila melanogaster reveals genes associated with Coxiella burnetii infection. Genetics, 217(3).

35. 10.1093/genetics/iyab005

36. Harbison, S. T., Kumar, S., Huang, W., McCoy, L. J., Smith, K. R., & Mackay, T. F. C. (2019). Genome-Wide Association Study of Circadian Behavior in Drosophila melanogaster. Behav Genet, 49(1), 60–82. 10.1007/s10519-018-9932-0

37. Harbison, S. T., McCoy, L. J., & Mackay, T. F. (2013). Genome-wide association study of sleep in Drosophila melanogaster. BMC Genomics, 14, 281. 10.1186/1471-2164-14-281

38. Harbison, S. T., Serrano Negron, Y. L., Hansen, N. F., & Lobell, A. S. (2017). Selection for long and short sleep duration in Drosophila melanogaster reveals the complex genetic network underlying natural variation in sleep. PLoS Genet, 13(e1007098), e10007098.

39. Hendricks, J. C., Finn, S. M., Panckeri, K. A., Chavkin, J., Williams, J. A., Sehgal, A., & Pack, A. I. (2000a). Rest in Drosophila is a sleep-like state. Neuron, 25(1), 129–138. 10.1016/s0896-6273(00)80877-6

40. Hendricks, J. C., Finn, S. M., Panckeri, K. A., Chavkin, J., Williams, J. A., Sehgal, A., & Pack, A. I. (2000b). Rest in Drosophila is a sleep-like state. Neuron, 25, 129–138.

41. Hori, A. (1986). Sleep characteristics in twins. Jpn J Psychiatry Neurol, 40(1), 35–46. 10.1111/j.1440-1819.1986.tb01610.x

42. Hu, Y., Flockhart, I., Vinayagam, A., Bergwitz, C., Berger, B., Perrimon, N., & Mohr, S. E. (2011). An integrative approach to ortholog prediction for disease-focused and other functional studies. BMC Bioinformatics, 12, 357. 10.1186/1471-2105-12-357

43. Huang, W., Massouras, A., Inoue, Y., Peiffer, J., Ramia, M., Tarone, A. M., Turlapati, L., Zichner, T., Zhu, D., Lyman, R. F., Magwire, M. M., Blankenburg, K., Carbone, M. A., Chang, K., Ellis, L. L., Fernandez, S., Han, Y., Highnam, G., Hjelmen, C. E.,…Mackay, T. F. (2014). Natural variation in genome architecture among 205 Drosophila melanogaster Genetic Reference Panel lines. Genome Res, 24(7), 1193–1208. 10.1101/gr.171546.113

44. Huber, R., Hill, S. L., Holladay, C., Biesiadecki, M., Tononi, G., & Cirelli, C. (2004). Sleep homeostasis in Drosophila melanogaster. Sleep, 27(4), 628–639. 10.1093/sleep/27.4.628

45. Hunter, C. M., Huang, W., Mackay, T. F., & Singh, N. D. (2016). The Genetic Architecture of Natural Variation in Recombination Rate in Drosophila melanogaster. PLoS Genet, 12(4), e1005951. 10.1371/journal.pgen.1005951

46. Iber, C. (2007). The AASM manual for the scoring of sleep and associated events: rules, terminology, and technical specification. *(No Title)*.

47. Jagannathan, S. R., Jeans, T., Van De Poll, M. N., & van Swinderen, B. (2024). Multivariate classification of multichannel long-term electrophysiology data identifies different sleep stages in fruit flies. Sci Adv, 10(8), eadj4399. 10.1126/sciadv.adj4399

48. Joyce, M., Falconio, F. A., Blackhurst, L., Prieto-Godino, L., French, A. S., & Gilestro, G. F. (2024). Divergent evolution of sleep in Drosophila species. Nat Commun, 15(1), 5091. 10.1038/s41467-024-49501-9

49. Kanehisa, M., Sato, Y., Kawashima, M., Furumichi, M., & Tanabe, M. (2016). KEGG as a reference resource for gene and protein annotation. Nucleic Acids Res, 44(D1), D457–462. 10.1093/nar/gkv1070

50. Keene, A. C., & Duboue, E. R. (2018). The origins and evolution of sleep. J Exp Biol, 221(Pt 11). 10.1242/jeb.159533

51. Keles, M. F., Sapci, A. O. B., Brody, C., Palmer, I., Mehta, A., Ahmadi, S., Le, C., Tastan, O., Keles, S., & Wu, M. N. (2025). FlyVISTA, an integrated machine learning platform for deep phenotyping of sleep in Drosophila. Sci Adv, 11(11), eadq8131. 10.1126/sciadv.adq8131

52. Koh, K., Joiner, W. J., Wu, M. N., Yue, Z., Smith, C. J., & Sehgal, A. (2008). Identification of SLEEPLESS, a sleep-promoting factor. Science, 321(5887), 372–376. 10.1126/science.1155942

53. Kuna, S. T., Maislin, G., Pack, F. M., Staley, B., Hachadoorian, R., Coccaro, E. F., & Pack, A. I. (2012). Heritability of performance deficit accumulation during acute sleep deprivation in twins. Sleep, 35(9), 1223–1233. 10.5665/sleep.2074

54. Li, C., Geng, C., Leung, H. T., Hong, Y. S., Strong, L. L., Schneuwly, S., & Pak, W. L. (1999). INAF, a protein required for transient receptor potential Ca(2+) channel function. Proc Natl Acad Sci U S A, 96(23), 13474–13479. 10.1073/pnas.96.23.13474

55. Li, Q., Jang, H., Lim, K. Y., Lessing, A., & Stavropoulos, N. (2021). insomniac links the development and function of a sleep-regulatory circuit. Elife, 10. 10.7554/eLife.65437

56. Linkowski, P., Kerkhofs, M., Hauspie, R., & Mendlewicz, J. (1991). Genetic determinants of EEG sleep: a study in twins living apart. Electroencephalogr Clin Neurophysiol, 79(2), 114–118. 10.1016/0013-4694(91)90048-9

57. Linkowski, P., Kerkhofs, M., Hauspie, R., Susanne, C., & Mendlewicz, J. (1989). EEG sleep patterns in man: a twin study. Electroencephalogr Clin Neurophysiol, 73(4), 279–284. 10.1016/0013-4694(89)90106-5

58. Lippert, C., Listgarten, J., Liu, Y., Kadie, C. M., Davidson, R. I., & Heckerman, D. (2011). FaST linear mixed models for genome-wide association studies. Nat Methods, 8(10), 833–835. 10.1038/nmeth.1681

59. Lobell, A. S., Kaspari, R. R., Serrano Negron, Y. L., & Harbison, S. T. (2017). The Genetic Architecture of Ovariole Number in Drosophila melanogaster: Genes with Major, Quantitative, and Pleiotropic Effects. G3(Bethesda), 7(7), 2391–2403. 10.1534/g3.117.042390

60. Loomis, A. L., Harvey, E. N., & Hobart, G. (1936). Electrical potentials of the human brain. Journal of Experimental Psychology, 19(3), 249–279. 10.1037/h0062089

61. Mackay, T. F., Richards, S., Stone, E. A., Barbadilla, A., Ayroles, J. F., Zhu, D., Casillas, S., Han, Y., Magwire, M. M., Cridland, J. M., Richardson, M. F., Anholt, R. R., Barron, M., Bess, C., Blankenburg, K. P., Carbone, M. A., Castellano, D., Chaboub, L., Duncan, L.,…Gibbs, R. A. (2012). The Drosophila melanogaster Genetic Reference Panel. Nature, 482(7384), 173–178. 10.1038/nature10811

62. Mackay, T. F. C., & Huang, W. (2018). Charting the genotype-phenotype map: lessons from the Drosophila melanogaster Genetic Reference Panel. Wiley Interdiscip Rev Dev Biol, 7(1). 10.1002/wdev.289

63. Mallon, E. B., Alghamdi, A., Holdbrook, R. T., & Rosato, E. (2014). Immune stimulation reduces sleep and memory ability in Drosophila melanogaster. PeerJ, 2, e434. 10.7717/peerj.434

64. Metaxakis, A., Tain, L. S., Gronke, S., Hendrich, O., Hinze, Y., Birras, U., & Partridge, L. (2014). Lowered insulin signalling ameliorates age-related sleep fragmentation in Drosophila. PLoS Biol, 12, e1001824.

65. Morey, M., Yee, S. K., Herman, T., Nern, A., Blanco, E., & Zipursky, S. L. (2008). Coordinate control of synaptic-layer specificity and rhodopsins in photoreceptor neurons. Nature, 456(7223), 795–799. 10.1038/nature07419

66. Moser, D., Anderer, P., Gruber, G., Parapatics, S., Loretz, E., Boeck, M., Kloesch, G., Heller, E., Schmidt, A., Danker-Hopfe, H., Saletu, B., Zeitlhofer, J., & Dorffner, G. (2009). Sleep classification according to AASM and Rechtschaffen & Kales: effects on sleep scoring parameters. Sleep, 32(2), 139–149. 10.1093/sleep/32.2.139

67. Nall, A. H., & Sehgal, A. (2013). Small-molecule screen in adult Drosophila identifies VMAT as a regulator of sleep. J Neurosci, 33(19), 8534–8540. 10.1523/JNEUROSCI.0253-13.2013

68. Neely, G. G., Hess, A., Costigan, M., Keene, A. C., Goulas, S., Langeslag, M., Griffin, R. S., Belfer, I., Dai, F., Smith, S. B., Diatchenko, L., Gupta, V., Xia, C. P., Amann, S., Kreitz, S., Heindl-Erdmann, C., Wolz, S., Ly, C. V., Arora, S.,…Penninger, J. M. (2010). A genome-wide Drosophila screen for heat nociception identifies alpha2delta3 as an evolutionarily conserved pain gene. Cell, 143(4), 628–638. 10.1016/j.cell.2010.09.047

69. Nitz, D. A., van Swinderen, B., Tononi, G., & Greenspan, R. J. (2002). Electrophysiological correlates of rest and activity in Drosophila melanogaster. Curr Biol, 12(22), 1934–1940. 10.1016/s0960-9822(02)01300-3

70. Oh, Y., Yoon, S. E., Zhang, Q., Chae, H. S., Daubnerova, I., Shafer, O. T., Choe, J., & Kim, Y. J. (2014). A homeostatic sleep-stabilizing pathway in Drosophila composed of the sex peptide receptor and its ligand, the myoinhibitory peptide. PLoS Biol, 12(10), e1001974. 10.1371/journal.pbio.1001974

71. Ozturk-Colak, A., Marygold, S. J., Antonazzo, G., Attrill, H., Goutte-Gattat, D., Jenkins, V. K., Matthews, B. B., Millburn, G., Dos Santos, G., Tabone, C. J., & FlyBase, C. (2024). FlyBase: updates to the Drosophila genes and genomes database. Genetics, 227(1). 10.1093/genetics/iyad211

72. Pfeiffenberger, C., & Allada, R. (2012). Cul3 and the BTB adaptor insomniac are key regulators of sleep homeostasis and a dopamine arousal pathway in Drosophila. PLoS Genet, 8(10), e1003003. 10.1371/journal.pgen.1003003

73. Pimentel, D., Donlea, J. M., Talbot, C. B., Song, S. M., Thurston, A. J. F., & Miesenbock, G. (2016). Operation of a homeostatic sleep switch. Nature, 536(7616), 333–337. 10.1038/nature19055

74. Qian, Y., Cao, Y., Deng, B., Yang, G., Li, J., Xu, R., Zhang, D., Huang, J., & Rao, Y. (2017). Sleep homeostasis regulated by 5HT2b receptor in a small subset of neurons in the dorsal fan-shaped body of drosophila. Elife, 6. 10.7554/eLife.26519

75. Qiu, Y., & Davis, R. L. (1993). Genetic dissection of the learning/memory gene dunce of Drosophila melanogaster. Genes Dev, 7(7B), 1447–1458. 10.1101/gad.7.7b.1447

76. Rabiner, L., & Juang, B. (1986). An introduction to hidden Markov models. IEEE ASSP Magazine, 3(1), 4–16. 10.1109/massp.1986.1165342

77. Rabiner, L. R. (1989). A tutorial on hidden Markov models and selected applications in speech recognition. Proceedings of the IEEE, 77(2), 257–286. 10.1109/5.18626

78. Raccuglia, D., Huang, S., Ender, A., Heim, M. M., Laber, D., Suarez-Grimalt, R., Liotta, A., Sigrist, S. J., Geiger, J. R. P., & Owald, D. (2019). Network-Specific Synchronization of Electrical Slow-Wave Oscillations Regulates Sleep Drive in Drosophila. Curr Biol, 29(21), 3611–3621 e3613. 10.1016/j.cub.2019.08.070

79. Rechtschaffen, A. (1968). A Manual of Standardized Terminology, Techniques and Scoring System for Sleep Stages of Human Subjects. Public Health Service.

80. Schwarz, G. (1978). Estimating the Dimension of a Model. The Annals of Statistics, 6(2). 10.1214/aos/1176344136

81. Seugnet, L., Suzuki, Y., Merlin, G., Gottschalk, L., Duntley, S. P., & Shaw, P. J. (2011). Notch signaling modulates sleep homeostasis and learning after sleep deprivation in Drosophila. Curr Biol, 21(10), 835–840. 10.1016/j.cub.2011.04.001

82. Shafer, O. T. (2025). 25 years of Drosophila “Sleep genes”. Fly, 19(1), 2502180. 10.1080/19336934.2025.2502180

83. Shannon, P., Markiel, A., Ozier, O., Baliga, N. S., Wang, J. T., Ramage, D., Amin, N., Schwikowski, B., & Ideker, T. (2003). Cytoscape: a software environment for integrated models of biomolecular interaction networks. Genome Res, 13(11), 2498–2504. 10.1101/gr.1239303

84. Shaw, P. J., Cirelli, C., Greenspan, R. J., & Tononi, G. (2000a). Correlates of sleep and waking in Drosophila melanogaster. Science, 287(5459), 1834–1837. 10.1126/science.287.5459.1834

85. Shaw, P. J., Cirelli, C., Greenspan, R. J., & Tononi, G. (2000b). Correlates of sleep and waking in Drosophila melanogaster. Science, 287, 1834–1837.

86. Shorter, J., Couch, C., Huang, W., Carbone, M. A., Peiffer, J., Anholt, R. R., & Mackay, T. F. (2015). Genetic architecture of natural variation in Drosophila melanogaster aggressive behavior. Proc Natl Acad Sci U S A, 112, E3555–E3563.

87. Singh, N. P., Ghosh, A., & Harbison, S. T. (2024). The Genetics of Sleep in Drosophila. In Genetics of Sleep and Sleep Disorders (pp. 7–56). 10.1007/978-3-031-62723-1_2

88. Spiers, J. G., Breda, C., Robinson, S., Giorgini, F., & Steinert, J. R. (2019). Drosophila Nrf2/Keap1 Mediated Redox Signaling Supports Synaptic Function and Longevity and Impacts on Circadian Activity. Front Mol Neurosci, 12, 86. 10.3389/fnmol.2019.00086

89. Stahl, B. A., Slocumb, M. E., Chaitin, H., DiAngelo, J. R., & Keene, A. C. (2017). Sleep-Dependent Modulation of Metabolic Rate in Drosophila. Sleep, 40(8). 10.1093/sleep/zsx084

90. Stavropoulos, N., & Young, M. W. (2011). insomniac and Cullin-3 regulate sleep and wakefulness in Drosophila. Neuron, 72(6), 964–976. 10.1016/j.neuron.2011.12.003

91. Szklarczyk, D., Kirsch, R., Koutrouli, M., Nastou, K., Mehryary, F., Hachilif, R., Gable, A. L., Fang, T., Doncheva, N. T., Pyysalo, S., Bork, P., Jensen, L. J., & von Mering, C. (2023). The STRING database in 2023: protein-protein association networks and functional enrichment analyses for any sequenced genome of interest. Nucleic Acids Res, 51(D1), D638–D646. 10.1093/nar/gkac1000

92. Tainton-Heap, L. A. L., Kirszenblat, L. C., Notaras, E. T., Grabowska, M. J., Jeans, R., Feng, K., Shaw, P. J., & van Swinderen, B. (2021). A Paradoxical Kind of Sleep in Drosophila melanogaster. Curr Biol, 31(3), 578–590 e576. 10.1016/j.cub.2020.10.081

93. Tea, J. S., Chihara, T., & Luo, L. (2010). Histone deacetylase Rpd3 regulates olfactory projection neuron dendrite targeting via the transcription factor Prospero. J Neurosci, 30(29), 9939–9946. 10.1523/JNEUROSCI.1643-10.2010

94. Tian, Y., Hu, W., Tong, H., & Han, J. (2012). Phototransduction in Drosophila. Sci China Life Sci, 55(1), 27–34. 10.1007/s11427-012-4272-4

95. Valatx, J. L., Bugat, R., & Jouvet, M. (1972). Genetic studies of sleep in mice. Nature, 238(5361), 226–227. 10.1038/238226a0

96. van Alphen, B., Semenza, E. R., Yap, M., van Swinderen, B., & Allada, R. (2021). A deep sleep stage in Drosophila with a functional role in waste clearance. Sci Adv, 7(4). 10.1126/sciadv.abc2999

97. van Alphen, B., Yap, M. H., Kirszenblat, L., Kottler, B., & van Swinderen, B. (2013). A dynamic deep sleep stage in Drosophila. J Neurosci, 33(16), 6917–6927. 10.1523/JNEUROSCI.0061-13.2013

98. Visser, I., & Speekenbrink, M. (2010). depmixS4: An R Package for Hidden Markov Models. Journal of Statistical Software, 36(7), 1–21. DOI 10.18637/jss.v036.i07

99. Viterbi, A. (1967). Error bounds for convolutional codes and an asymptotically optimum decoding algorithm. IEEE Transactions on Information Theory, 13(2), 260–269. 10.1109/TIT.1967.1054010 , ISSN= 1557-9654

100. Vonesch, S. C., Lamparter, D., Mackay, T. F., Bergmann, S., & Hafen, E. (2016). Genome-Wide Analysis Reveals Novel Regulators of Growth in Drosophila melanogaster. PLoS Genet, 12(1), e1005616. 10.1371/journal.pgen.1005616

101. Webb, W. B., & Campbell, S. S. (1983). Relationships in sleep characteristics of identical and fraternal twins. Arch Gen Psychiatry, 40(10), 1093–1095. 10.1001/archpsyc.1983.01790090055008

102. Wiggin, T. D., Goodwin, P. R., Donelson, N. C., Liu, C., Trinh, K., Sanyal, S., & Griffith, L. C. (2020). Covert sleep-related biological processes are revealed by probabilistic analysis in Drosophila. Proc Natl Acad Sci U S A, 117(18), 10024–10034. 10.1073/pnas.1917573117

103. Wu, K. J., Kumar, S., Serrano Negron, Y. L., & Harbison, S. T. (2018). Genotype influences day-to-day variability in sleep in Drosophila melanogaster. *Sleep*, zsx205, 10.1093/sleep/zs1205.

104. Wu, M., Robinson, J. E., & Joiner, W. J. (2014). SLEEPLESS is a bifunctional regulator of excitability and cholinergic synaptic transmission. Curr Biol, 24(6), 621–629. 10.1016/j.cub.2014.02.026

105. Wu, M. N., Joiner, W. J., Dean, T., Yue, Z., Smith, C. J., Chen, D., Hoshi, T., Sehgal, A., & Koh, K. (2010). SLEEPLESS, a Ly-6/neurotoxin family member, regulates the levels, localization and activity of Shaker. Nat Neurosci, 13(1), 69–75. 10.1038/nn.2454

106. Xia, S., & Chiang, A. S. (2009). NMDA Receptors in Drosophila. In A. M. Van Dongen (Ed.), Biology of the NMDA Receptor.

107. Xia, S., Miyashita, T., Fu, T. F., Lin, W. Y., Wu, C. L., Pyzocha, L., Lin, I. R., Saitoe, M., Tully, T., & Chiang, A. S. (2005). NMDA receptors mediate olfactory learning and memory in Drosophila. Curr Biol, 15(7), 603–615. 10.1016/j.cub.2005.02.059

108. Yap, M. H. W., Grabowska, M. J., Rohrscheib, C., Jeans, R., Troup, M., Paulk, A. C., van Alphen, B., Shaw, P. J., & van Swinderen, B. (2017). Oscillatory brain activity in spontaneous and induced sleep stages in flies. Nat Commun, 8(1), 1815. 10.1038/s41467-017-02024-y

109. Yin, L., Zhang, H., Tang, Z., Xu, J., Yin, D., Zhang, Z., Yuan, X., Zhu, M., Zhao, S., Li, X., & Liu, X. (2021). rMVP: A Memory-efficient, Visualization-enhanced, and Parallel-accelerated Tool for Genome-wide Association Study. Genomics Proteomics Bioinformatics, 19(4), 619–628. 10.1016/j.gpb.2020.10.007

110. Zhang, S., Ross, K. D., Seidner, G. A., Gorman, M. R., Poon, T. H., Wang, X., Keithley, E. M., Lee, P. N., Martindale, M. Q., Joiner, W. J., & Hamilton, B. A. (2015). Nmf9 Encodes a Highly Conserved Protein Important to Neurological Function in Mice and Flies. PLoS Genet, 11(7), e1005344. 10.1371/journal.pgen.1005344

111. Zwarts, L., Vanden Broeck, L., Cappuyns, E., Ayroles, J. F., Magwire, M. M., Vulsteke, V., Clements, J., Mackay, T. F., & Callaerts, P. (2015). The genetic basis of natural variation in mushroom body size in Drosophila melanogaster. Nat Commun, 6, 10115. 10.1038/ncomms10115

